# Lipidomic alterations in the cerebral cortex and white matter in sporadic Alzheimer’s disease

**DOI:** 10.1101/2022.11.04.515175

**Authors:** Elia Obis, Joaquim Sol, Pol Andres-Benito, Meritxell Martín-Gari, Natàlia Mota-Martorell, José Daniel Galo-Licona, Gerard Piñol-Ripoll, Manuel Portero-Otin, Isidro Ferrer, Mariona Jové, Reinald Pamplona

## Abstract

**Aims:** Non-targeted lipidomics analysis was conducted in post-mortem human frontal cortex area 8 (GM) and white matter of the frontal lobe *centrum semi-ovale* (WM) to identify lipidomes in middle-aged individuals with no neurofibrillary tangles and senile plaques, and cases at progressive stages of sporadic Alzheimer’s disease (sAD).

**Methods:** Lipidomic analysis using an LC-MS/MS platform was carried out in selected cases with suitable post-mortem lacking co-morbidities and concomitant brain pathologies. Complementary data were obtained using RT-qPCR and immunohistochemistry.

**Results:** The WM shows an adaptive lipid phenotype resistant to lipid peroxidation, characterized by a lower fatty acid unsaturation, peroxidizability index, and higher ether lipid content than the GM. Changes in the lipidomic profile more marked in the WM than in GM occur in AD with disease progression. WM alterations are characterized by a decline in the content of branched fatty acid esters of hydroxy fatty acids (FAHFA), diacylglycerols (DG), triacylglycerols (TG), glycerophospholipids (GP) (especially ether lipids), and sphingolipids (especially sulfatides, ceramides, and glycosphingolipids). The GM acquires a fatty acid profile prone to peroxidation in sAD, while WM reinforces its peroxidation-resistant trait. Transcriptomic data point to additional defects of peroxisomal function.

**Conclusions:** Four functional categories are associated with the different lipid classes affected in sAD: membrane structural composition, bioenergetics, antioxidant protection, and bioactive lipids, with deleterious consequences affecting both neurons and glial cells favoring disease progression.

## 1. Introduction

The cerebral white matter (WM) in brain aging shows a reduction of the total volume and progressive alteration of the structural integrity, manifested as diffuse myelin decrease and focal lesions that lead to impaired brain connectivity (Hachinski et al., 1987; Scheltens et al., 1995; Barber et al., 1999; Mamer et al., 2003; Holland et al., 2008; Salat et al., 2009; Gao et al., 2011; Erten-Lyons et al., 2013; Wharton et al., 2015; Liu et al., 2017). These alterations bring to a cognitive and neuropsychiatric detriment (Bartzokis et al., 2003; Bartzokis et al., 2004; Bennett et al., 2014; Kavroulakis et al., 2018). Modification in the number and characteristics of oligodendrocytes and oligodendroglial precursor cells, responsible of myelin homeostasis, occurs in aged non-human primates and humans (Kohama et al., 2012).

Neuroimaging methods also reveal alterations in the WM in sporadic Alzheimer’s disease (sAD), including reduced WM size, hyper-intensities, myelin and axon damage, and reduced connectivity, in association with cognitive impairment (Scheltens et al., 1995; Barber et al., 1999; Holland et al., 2008; Bartzokis et al., 2003; Diaz et al., 1991; Rose et al., 2000; Medina et al., 2006; Wang et al., 2011; Selnes et al., 2012; Radanovic et al., 2013; Brickman, 2013; Molinuevo et al., 2014; Hoy et al., 2017; Bouhrara et al., 2018; Joki et al., 2018). WM alterations precede the appearance of clinical symptoms (Lee et al., 2016), and WM further deteriorates with AD progression (Medina et al., 2006; Selnes et al., 2012; Molinuevo et al., 2014; Hoy et al., 2017; Bouhrara et al., 2018). The cerebral WM’s atrophy and demyelination are also pinpointed in post-mortem neuropathological studies (de la Monte, 1989; Gouw et al., 2008; Ihara et al., 2010; Ferrer and Andres-Benito, 2020). Therefore, loss of WM integrity is considered a key component of sAD, thus contributing to neural disconnection, cognitive impairment, and dementia (Bartzokis et al., 2004).

Lipids have favored brain evolution to attain its structural and functional complexity (Crawford et al., 2001; Broadhurst et al., 2002; Crawford et al., 2014). The human brain is one of the tissues richest in lipid content (Sastry, 1985), with the most extensive diversity of lipid classes (for instance, glycerolipids (GLs), glycerophospholipids (GPs), sphingolipids (SLs) and cholesterol), and lipid molecular species. The human brain also has a wide diversity of functional properties covering the structural and functional integrity of neuronal and glial cell membranes, generation of lipid mediators, and chemical reactivity of the acyl chains (Naudi et al., 2015). Adult human brain lipids undergo slow but progressive and significant region-specific modifications in their concentration and distribution during aging. Thus, total lipid content ―including fatty acids, GPs (especially ether lipids), SLs, and cholesterol― decreases after age 50 (Naudi et al., 2015). Lipoxidation-derived protein damage also increases with age in a region-specific manner (Horrocks et al., 1981; Soderberg et al., 1990; Svennerholm et al., 1991; Svennerholm et al., 1994; McNamara et al., 2008; Ledesma et al., 2012; Hancock et al., 2015; Norris et al., 2015; Dominguez et al., 2016; Cabre et al., 2017; Hancock et al., 2017; Cabre et al., 2018; Diaz et al., 2018; Dominguez-Gonzalez et al., 2018; Jove et al., 2019; Pamplona et al., 2019; Jove et al., 2021; Mota-Martorell et al., 2022).

Brain lipidomic analyses in sAD have identified multiple disease-specific lipid alterations and lipid-derived molecular damage (Naudi et al., 2015; Farooqui et al., 1988; Pamplona et al., 2005; Han, 2010; Haughey et al., 2010; Martin et al., 2010; Terni et al., 2010; Frisardi et al., 2011; Wood et al., 2012; Zhu et al., 2012; Kosicek & Hecimovic, 2013; Sultana et al., 2013; Touboul & Gaudin, 2014; Zabel et al., 2018; Jove et al., 2021). These alterations include depletion of ether lipids and sulfatides, increased ceramides (Cer), and severe lipoxidative damage. Lipidomic modifications detected at the early stages of sAD aggravate the disease’s progression. Nevertheless, described changes in sAD mainly refer to different regions of the grey matter, whereas studies of lipid changes in the WM in sAD are minimal (Wood et al., 2015).

A seminal study described differences in the lipid composition between human WM and grey matter through the lifespan (O’Brien and Sampson, 1965). In the adult brain, the total amount of lipids in the grey matter was 36-40% and 19-66% in the WM. The WM showed higher levels of sphingolipids (including sphingomyelins (SMs), cerebrosides, cerebrosides sulfatides, Cer, and cholesterol in comparison with the grey matter. No age-related WM changes were observed in total GPs, but glycerophosphatidylserines (PS) were increased, and glycerophosphatidylcholines (PCs) decreased (O’Brien and Sampson, 1965).

This study was designed to assess lipid alterations separately in the frontal cortex area 8 and WM of the frontal lobe’s *centrum semi-ovale* in aging and sAD at different stages of progression. Cases with clinical and pathological co-morbidities were not included in the study. LC-MS/MS platform and gas chromatography was used for the lipidomics study. mRNA expression and proteins involved in lipid metabolism were analyzed by RT-qPCR and immunohistochemistry, respectively. We aimed to identify lipidome differences between the cerebral cortex and WM in brain aging and sAD using novel high throughput mass spectrometry-based techniques combined with protein expression analysis involved in selected lipid metabolism pathways, demonstrating that sAD is associated with altered lipidome profiles.

## 2. Material and Methods

### 2.1. Selection of human samples

Post-mortem samples of fresh-frozen tissue from the frontal cortex area 8 and *centrum semi-ovale* of the frontal lobe were obtained from the Institute of Neuropathology HUB-ICO-IDIBELL Biobank, following the guidelines of Spanish legislation on this matter (Real Decreto 1716/2011), and the approval of the local ethics committee.

One hemisphere was immediately cut in coronal sections 1□cm thick and selected areas of the encephalon were rapidly dissected, frozen on metal plates over dry ice, placed in individual air-tight plastic bags, and stored at -80°C until used for biochemical studies. The other hemisphere was fixed by immersion in 4% buffered formalin for three weeks for morphological studies. The neuropathological study was carried out on selected 4-μm-thick de-waxed paraffin sections of 20 representative regions. Sections were stained with hematoxylin and eosin, periodic acid-Schiff (PAS), and Klüver-Barrera, or processed for immunohistochemistry for β-amyloid, phospho-tau (clone AT8), α-synuclein, αB-crystallin, TDP-43, ubiquitin, p62, glial fibrillary acidic protein, CD68, and Iba1 (Ferrer, 2015). The post-mortem delay varied from 1□hour and 30□minutes to 16□hours (**Table 1**). The brain pH at the autopsy was between 6.2 and 6.4, and the RNA integrity number (RIN) was higher than 6, thus ensuring the biological sample’s quality (Ferrer et al., 2007; Ferrer, 2008; Ferrer, 2015).

**Table 1:** Summary of middle-aged individuals without NFTs and SPs in any brain region (MA) and cases at different Braak stages of Alzheimer’s disease (AD) without co-morbidities and concomitant brain pathologies.

| Case ID | Diagnosis | Sex | Age | PM delay | RIN WM |
| --- | --- | --- | --- | --- | --- |
| 1 | AD I/0 | M | 56 | 07 h 10 min | 7.80 |
| 2 | AD I/A | W | 74 | 02 h 45 min | 7.70 |
| 3 | AD I/A | W | 57 | 05 h 00 min | 6.50 |
| 4 | AD I/A | M | 66 | 09 h 45 min | 6.30 |
| 5 | AD II/0 | M | 67 | 07 h 15 min | 6.90 |
| 6 | AD II/A | W | 88 | 08 h 00 min | 6.90 |
| 7 | AD II/A | W | 86 | 02 h 15 min | 8.30 |
| 8 | AD III/0 | M | 81 | 01 h 30 min | 7.60 |
| 9 | AD III/A | W | 82 | 02 h 00 min | 7.20 |
| 10 | AD III/A | W | 77 | 03 h 10 min | 6.40 |
| 11 | AD III/B | W | 76 | 03 h 50 min | 7.20 |
| 12 | AD IV/A | W | 80 | 02 h 45 min | 5.40 |
| 13 | AD V/B | W | 74 | 05 h 30 min | 7.70 |
| 14 | AD V/B | M | 86 | 04 h 15 min | 7.90 |
| 15 | AD V/B | M | 73 | 04 h 30 min | 6.90 |
| 16 | AD VI/C | W | 72 | 09 h 30 min | 6.40 |
| 17 | AD V/C | M | 85 | 03 h 45 min | 7.80 |
| 18 | AD V/C | M | 77 | 16 h 00 min | 7.10 |
| 19 | MA | W | 62 | 11 h 00 min | 8.40 |
| 20 | MA | W | 53 | 03 h 00 min | 7.70 |
| 21 | MA | M | 39 | 09 h 15 min | 7.10 |
| 22 | MA | M | 57 | 03 h 00 min | 8.20 |
| 23 | MA | M | 50 | 17 h 15 min | 8.00 |
| 24 | MA | W | 54 | 06 h 45 min | 7.20 |

M: male; F: female; PM: post-mortem delay; RIN: RNA integrity numbers; WM: white matter of the frontal lobe *centrum semi-ovale*.

sAD cases were categorized according to Braak and Braak (1991) neurofibrillary tangle (NFT) and β-amyloid stages as ADI–II/0-A (n = 7, men: 3, women: 4); ADIII– IV/0-C (n = 5, men: 1, women: 4), and ADV–VI/B-C (n = 6, men: 4, women: 2) (**Table 1**). Cases with concomitant pathologies and co-morbidities were excluded, including age-related neurodegenerative diseases (tauopathies, Lewy body diseases, TDP-43 proteinopathy), hippocampal sclerosis; and those who had suffered from cerebrovascular disease, arterial hypertension, type II diabetes, hyperlipidemia, cardiac, hepatic, renal failure, and respiratory insufficiency. All selected cases were sporadic; familial AD was not included in this study. AD cases at stages I–II/0-A had no neurological symptoms; AD cases at stages III–IV/0-C had no neurological symptoms or were affected by mild cognitive impairment; AD cases at stages V–VI/B-C had severe cognitive impairment or dementia.

The last group (n = 6) comprised middle-aged cases (MA) of three men and three women. MA did not have clinical risk factors and co-morbidities mentioned in previous paragraphs; they did not have neurological or mental diseases, and the neuropathological study did not show abnormalities. This MA group must not be interpreted as an age-matched control group but as a control group of normal WM in MA individuals. The total number of MA and AD cases in this series is detailed in **Table 1**.

### 2.2. Fatty acid profiling

Fatty acyl groups were analyzed as methyl esters derivatives by gas chromatography as previously described (Mota-Martorell et al., 2022). For tissue homogenization, 50mg of cerebral cortex and WM was processed in a buffer containing 180mM KCl, 5mM MOPS, 2mM EDTA, 1mM diethylenetriaminepentaacetic acid, and 1μM butylated hydroxytoluene. Tissue samples were randomized prior to lipid extraction. Quality control samples were included at a ratio of 1:5. Total lipids from samples were extracted into chloroform:methanol (2:1, v/v) in the presence of 0.01% (w/v) butylated hydroxytoluene. The chloroform phase was evaporated under nitrogen, and the fatty acyl groups were transesterified by incubation in 2.5mL of 5% (v/v) methanolic HCl at 75°C for 90 min. The resulting fatty acid methyl esters were extracted by adding 1mL of saturated NaCl solution and 2.5mL of n-pentane. The n-pentane phase was separated and evaporated under N_2_. The residue was dissolved in 50µL of CS_2_, and 2µL was used for analysis. Separation was performed by a DBWAX capillary column (30m x 0.25mm x 0.20μm) in a GC System 7890A with a Series Injector 7683B and an FID detector (Agilent Technologies, Barcelona, Spain). The sample injection was in splitless mode. The injection port was maintained at 250°C, and the detector at 250°C. The program consisted of 5 min at 145°C, followed by 2°C/min to 245°C, and finally 245°C for 10 min, with a post-run at 250°C for 10 minutes. The total run time was 65 minutes, with a post-run time of 10 minutes. Identification of fatty acid methyl esters was made by comparison with authentic standards (Larodan Fine Chemicals, Malmö, Sweden) using specific software of data analysis for GC from Agilent (OpenLAB CDS ChemStation v. C.01.10; Agilent Technologies, Barcelona, Spain) and subsequent expert’s revision and confirmation. Results are expressed as mol%.

The following fatty acyl indices were also calculated: saturated fatty acids (SFA); unsaturated fatty acids (UFA); monounsaturated fatty acids (MUFA); polyunsaturated fatty acids (PUFA) from n-3 and n-6 series (PUFAn-3 and PUFAn-6, respectively); and average chain length, ACL = [(Σ%Total_14_ x 14) + (Σ% Total_16_×16) + (Σ%Total_18_×18) + (Σ%Total_20_×20) + (Σ% Total_22_×22) + (Σ% Total_24_×24)]/100. The density of double-bonds in the membrane was calculated with the Double-Bond Index, DBI = [(1 × Σmol% monoenoic) + (2 × Σmol% dienoic) + (3 × Σmol% trienoic) + (4 × Σmol% tetraenoic) + (5 × Σmol% pentaenoic) + (6 × Σmol% hexaenoic)]. Membrane susceptibility to peroxidation was calculated with the Peroxidizability Index, PI (a) = [(0.025 × Σmol% monoenoic) + (1 × Σmol% dienoic) + (2 × Σmol% trienoic) + (4 × Σmol% tetraenoic) + (6 × Σmol% pentaenoic) + (8 × Σmol% hexaenoic)] (Mota-Martorell et al., 2022), and PI (b) = [(0.015 × Σmol% monoenoic) + (1 × Σmol% dienoic) + (2 × Σmol% trienoic) + (3 × Σmol% tetraenoic) + (4 × Σmol% pentaenoic) + (5 × Σmol% hexaenoic)] (Yin et al., 2011).

Elongase and desaturase activities were estimated from specific product/substrate ratios (Guillou et al., 2010): Elovl3(n-9) = 20:1n-9/18:1n-9; Elovl6 = 18:0/16:0; Elovl1-3-7a = 20:0/18:0; Elovl1-3-7b = 22:0/20:0; Elovl1-3-7c = 24:0/22:0; Elovl5(n-6) = 20:2n-6/18:2n-6; Elovl2-5 (n-6) = 22:4n-6/20:4n-6; Elovl 2-5(n-3) = 22:5n-3/20:5n-3, Elovl 2(n-3)= 24:5n-3/22:5n-3, Δ9(n-7) = 16:1n-7/16:0; Δ9(n-9) = 18:1n-9/18:0; Δ5(n-6) = 20:4n-6/20:3n-6; Δ6(n-3) = 18:4n-3/18:3n-3; and Δ6(n-3) = 24:6n-3/24:5n-3. Finally, the peroxisomal β-oxidation was estimated according to the 22:6n-3/24:6n-3 ratio.

### 2.3. Non-targeted lipidomic analysis

#### Sample preparation

For the lipid extraction, 10μL of the homogenized tissue were mixed with 5μL of MiliQ water and 20μL of ice-cold methanol. Samples were vigorously shaken by vortexing for 2 min, and then 250μL of methyl tert-butyl ether (MTBE), containing internal lipid standards (see Supplementary **Table S1**), were added. Samples were immersed in a water bath (ATU Ultrasonidos, Valencia, Spain) with an ultrasound frequency and power of 40 kHz and 100 W, respectively, at 10°C for 30 min. Then, 25μL MiliQ water was added to the mixture, and the organic phase was separated by centrifugation (1,400 g) at 10°C for 10 min (Pizarro et al., 2013). Lipid extracts in the upper phase were subjected to mass spectrometry. A pool of all lipid extracts was prepared and used as quality control. Internal isotopically labeled lipid standards for each class were used for signal normalization (Pradas et al., 2018). Stock solutions were prepared by dissolving lipid standards in MTBE at a concentration of 1mg/mL, and working solutions were diluted to 2.5μg/mL in MTBE.

#### LC-MS analysis

Lipid extracts were analyzed following a previously published method (Castro-Perez et al., 2010). Lipid extracts were subjected to liquid chromatography-mass spectrometry using a UPLC 1290 series coupled to ESI-Q-TOF MS/MS 6545 (Agilent Technologies, Barcelona, Spain). The sample compartment of the UHPLC was refrigerated at 4°C, and for each sample, 10μL of lipid extract was applied onto a 1.8μm particle 100 × 2.1mm id Waters Acquity HSS T3 column (Waters, Milford, MA, USA) heated at 55°C. The flow rate was 400μL/min with solvent A composed of 10mM ammonium acetate in acetonitrile-water (40:60, v/v) and solvent B composed of 10mM ammonium acetate in acetonitrile-isopropanol (10:90, v/v). The gradient started at 40% of mobile phase B, reached 100% B in 10 min, and held for 2 min. Finally, the system was switched back to 60% of mobile phase B and was equilibrated for 3 min. Duplicate runs of the samples were performed to collect positive and negative electrospray-ionized lipid species in a TOF mode, operated in full-scan mode at 100 to 3000 m/z in an extended dynamic range (2GHz), using N_2_ as nebulizer gas (5L/min, 350°C). The capillary voltage was set at 3500V with a scan rate of one scan/s. Continuous infusion using a double spray with masses 121.050873, 922.009798 (positive ion mode) and 119.036320, 966.000725 (negative ion mode) was used for in-run calibration of the mass spectrometer (Pradas et al., 2019).

#### Lipidomic Data analysis and Annotation

MassHunter Qualitative Analysis Software (Agilent Technologies, Barcelona, Spain) was used to obtain the molecular features of the samples, representing different co-migrating ionic species of a given molecular entity using the Molecular Feature Extractor algorithm (Agilent Technologies, Barcelona, Spain). MassHunter Mass Profiler Professional Software (Agilent Technologies, Barcelona, Spain) and Metabolanalyst Software (Xia and Wishart, 2016; Chong and Xia, 2018) were used to perform a non-targeted lipidomic analysis of the obtained data. Only those features with a minimum of 2 ions were selected. After that, the molecular characteristics in the samples were aligned using a retention time window of 0.1 % ± 0.25 min and 30.0 ppm ± 2.0mDa. Only features found in at least 70% of the QC samples accounted for the correction of individual bias, and the signal was corrected using a LOESS approach (Dunn et al., 2011; Broadhurst et al., 2018). Multivariate statistics (Principal Component Analysis (PCA), Partial Least-squares Discriminant Analysis (PLS-DA), and Hierarchical and Classification Analyses) were performed using Metaboanalyst software. For annotation features representing significant differences by Univariate statistics ANOVA (p < 0.05), defined by exact mass and retention time, were searched against the HMDB (Wishart et al., 2018) (accuracy < 30ppm) and LIPID MAPS (Fahy et al., 2007) databases (accuracy < 20ppm). The identities obtained were compared to the authentic standards’ retention times. Finally, identities were confirmed by searching experimental MS/MS spectra against *in silico* libraries, using HMDB and LipidMatch, an R-based tool for lipid identification (Koelmel et al., 2017).

### 2.4. Immunohistochemistry

Formalin-fixed, paraffin-embedded, de-waxed sections 4 µm thick of the frontal cortex in five MA cases were processed for specific immunohistochemistry. The sections were boiled in citrate buffer (20 min) to retrieve protein antigenicity. Endogenous peroxidases were blocked by incubation in 10% methanol-1% H_2_O_2_ solution (15 min) followed by 3% normal horse serum solution. Then the sections were incubated at 4ºC overnight with one of the primary rabbit polyclonal antibodies: ACAA1, 3-ketoacyl-CoA thiolase (MyBioSource MBS1492126) used at a dilution of 1/100; FAS, Fatty Acid Synthase (C20G5, Cell Signaling 3180) diluted 1/50, and SCD, Stearoyl-CoA desaturase (MyBioSource, BS421254) used at a dilution of 1/50. After incubation with the primary antibody, the sections were incubated with EnVision+ system peroxidase (Dako, Agilent Technologies, Santa Clara, CA, USA) for 30 min at room temperature. The peroxidase reaction was visualized with diaminobenzidine and H_2_O_2_. Control of the immunostaining included omission of the primary antibody; no signal was obtained following incubation with only the secondary antibody. Sections were slightly counterstained with hematoxylin. Due to the individual variability of the immunostaining, no attempt at quantification was performed; immunohistochemistry was used to assess the localization of the enzymes.

### 2.5. RNA extraction and RT-qPCR validation

RNA from frozen FC area 8 was extracted following the supplier’s instructions (RNeasy Mini Kit, Qiagen® GmbH, Hilden, Germany). RNA integrity and 28S/18S ratios were determined with the Agilent Bioanalyzer (Agilent Technologies Inc, Santa Clara, CA, USA) to assess RNA quality, and the RNA concentration was evaluated using a NanoDrop™ Spectrophotometer (Thermo Fisher Scientific). Complementary DNA (cDNA) preparation used a High-Capacity cDNA Reverse Transcription kit (Applied Biosystems, Foster City, CA, USA) following the protocol provided by the supplier. TaqMan RT-qPCR assays were duplicated for each gene on cDNA samples in 384-well optical plates using an ABI Prism 7900 Sequence Detection system (Applied Biosystems, Life Technologies, Waltham, MA, USA). For each 10μL TaqMan reaction, 2.25μL cDNA was mixed with 0.25μL 20x TaqMan Gene Expression Assays and 2.50μL of 2x TaqMan Universal PCR Master Mix (Applied Biosystems). The identification numbers and names of TaqMan probes are shown in **Supplementary Table S2**. Values of β-glucuronidase (*GUS-*β mRNA) were used as internal controls for normalization. The parameters of the reactions were 50°C for 2 min, 95°C for 10 min, and 40 cycles of 95°C for 15 sec and 60°C for 1 min. Finally, Sequence Detection Software (SDS version 2.2.2, Applied Biosystems) was used to capture TaqMan PCR data. The data were analyzed with the double-delta cycle threshold (ΔΔCT) method.

### 2.6. Statistics

For fatty acids, all values were expressed as means ± standard error of the mean (SEM). Comparisons between groups were analyzed by ANOVA followed by DMS tests for paired groups. The minimum level of statistical significance was set at p < 0.05 in all the analyses. For lipidomics, multivariate and univariate statistics were calculated using Metaboanalyst (Pang et al., 2020). Spearman correlations were performed with R version 4.0.3 (R Core Team (2019). R: A language and environment for statistical computing; R Foundation for Statistical Computing, Vienna, Austria. URL https://www.R-project.org/.), and p-values were adjusted with Benjamini-Hochberg’s false discovery rate (FDR). For transcriptomics, the statistical study was performed using the T-student test or ANOVA when necessary. The significance level was set at * p < 0.05, ** p < 0.01 and *** p < 0.001 vs MA; # p < 0.05 and ## p < 0.01 vs AD I-II/0-A; and $ p < 0.05 vs AD III-IV/A-B.

## 3. Results

### 3.1. Lipidomic profiles differ in cerebral cortex and WM

The first goal of the present study was to characterize the potential differences between the grey matter of the frontal cortex (GM) and WM of the frontal lobe’s *centrum semi-ovale* in healthy adult individuals according to their lipid composition. Firstly, we applied an untargeted lipidomic methodology to obtain global information about the differences between GM and WM lipidomes. Then, we characterized the fatty acid profiles of lipidomes from both GM and WM using a targeted approach.

Baseline correction, peak picking, and peak alignment were performed on untargeted approach acquired data resulting in a total of 9,207 molecules from negative and positive ionization modes. After quality control assessment, filtering, and correcting the signal, 2,048 features were used for multivariate and univariate statistical analysis. Multivariate statistics (**Figure 1A-D**) revealed marked differences in lipidome profiles, indicating that GM and WM show region-specific lipidomes. These results were reinforced by the detection, characterization, and quantification of 24 different fatty acids (**Figure 1E-H**).

**Figure 1:**
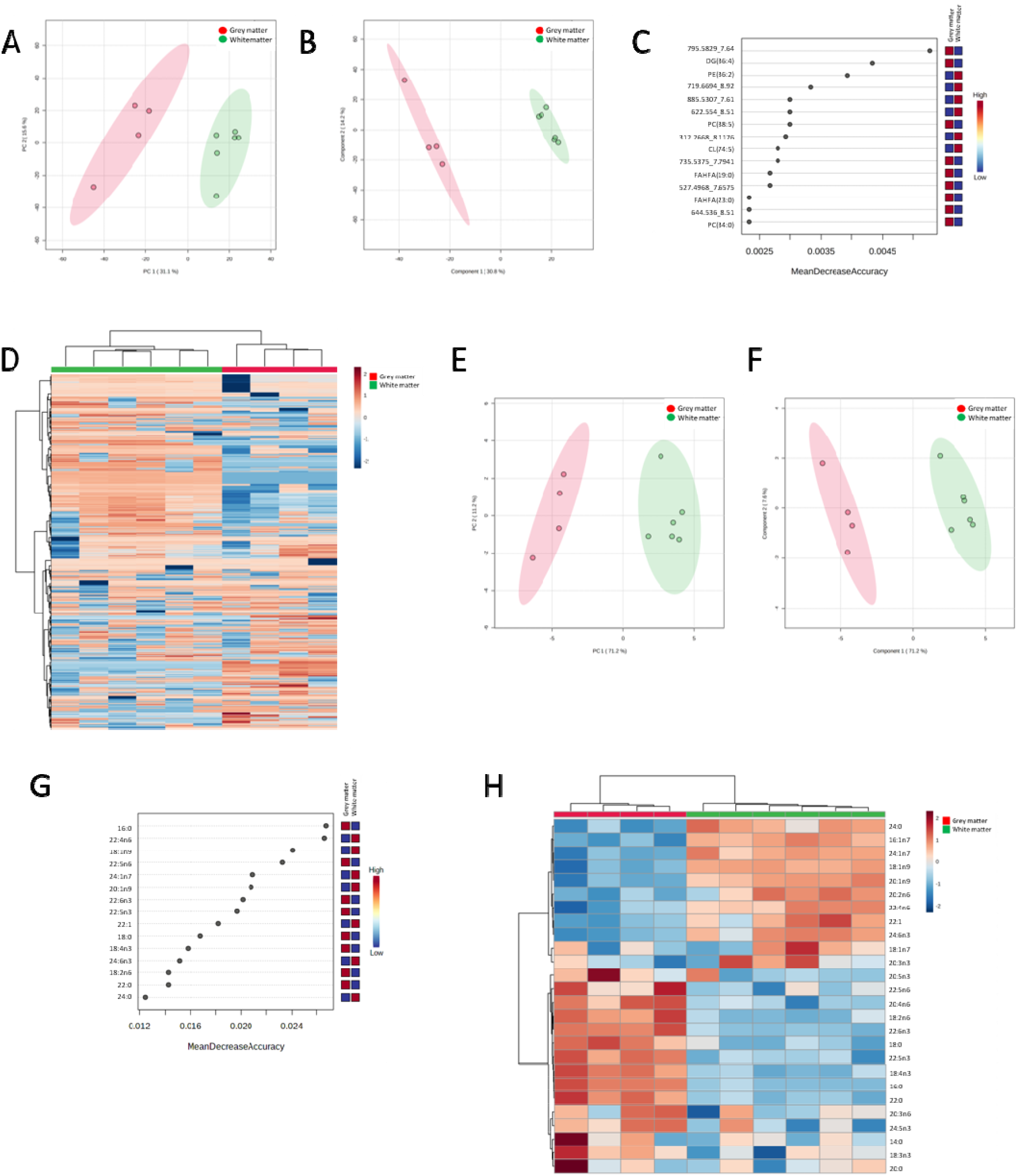
Untargeted and targeted lipid profiles distinguish grey and white matter brain tissues. A) Principal component analysis (PCA) scores plot of samples whole lipidome. B) Partial least squares discriminant analysis (PLS-DA) scores plot of samples lipidome. Leave One Out Cross-Validation (LOOCV) accuracy: 1.0, R2: 0.97, and Q2: 0.81 (one component). C) Random Forest classification of samples VIP plot according to Mean Decrease in Accuracy using brain white and grey matter whole lipidome. OBB Error: 0.0. D) Heatmap clustering analysis of samples whole lipidome. E) Principal component analysis (PCA) scores plot of samples fatty acids (FA) profile. F) Partial least squares discriminant analysis (PLS-DA) scores plot of samples FA profile. Leave One Out Cross-Validation (LOOCV) accuracy: 1.0, R2: 0.96, and Q2: 0.93 (one component). G) Random Forest classification of samples VIP plot according to Mean Decrease in Accuracy using brain white and grey matter FA profile. OBB Error: 0.0. H) Heatmap clustering analysis of samples fatty acids (FA) profile.

Univariate statistics of untargeted lipidomics revealed 270 statistically different lipid species between GM and WM (adjusted p-value FDR <0.07) (Supplementary **Table S3** and **Table S4**). We annotated 237 lipids (based on exact mass, retention time, isotopic distribution, and MS/MS spectrum) (Supplementary **Table S3**). Identified lipids belong to five major categories (**Figure 2**): i) Fatty acyls, covering seven esters of fatty acids, as fatty acid esters of hydroxyl fatty acids (FAHFA), and one AcylCoAs, the Retinoyl-CoA; ii) Glycerolipids, including 15 diacylglycerols (DG) and 14 triacylglycerols (TG); iii) Glycerophospholipids (GP), including 42 PCs, 58 glycerophosphoethanolamines (PEs), seven glycerophosphoglycerols (PGs), three cardiolipins (CLs), 14 PS, and four phosphatidic acids (PAs); iv) Sphingolipids, counting 25 Cer, one ceramide phosphate (CerP), 26 glycosylceramides (HexCer), nine sulfatides, and 10 SM; and v) Sterol lipids: two cholesteryl esters (CEs). The remaining thirty-three were unknown lipid species (Supplementary **Table S4**). The WM, in comparison with GM, exhibited lower concentration of DG, PA, and CE, and significant enrichment in TG(O), PC, PE, sulfatides, Cer, glycosphingolipids, and SM (Supplementary **Table S3**). Remarkably, several lipid species showing enrichment in WM corresponded to alkyl and alkenyl ethers, mostly presented as TG, PC, and PE species. For FAHFA, TG, PG, CL, and PS, the lower or enriched condition in WM is lipid species-dependent (Supplementary **Table S3**).

**Figure 2:**
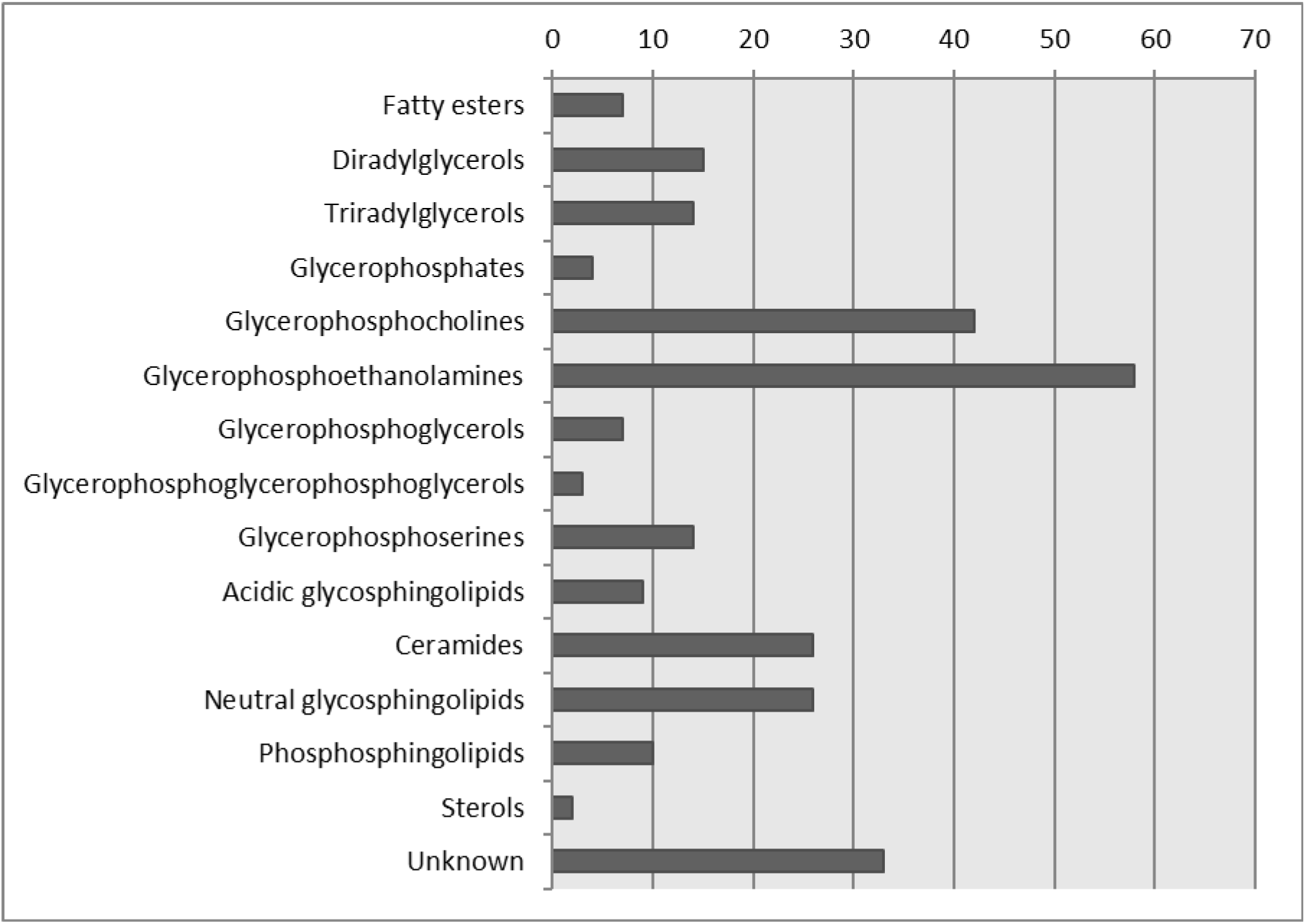
Lipid classes and numbers of identified lipid molecular species differ in WM and GM. The 237 identified lipid features were named and organized into five major categories, according to the LipidMaps database. The first category, Fatty acyls, comprised esters of fatty acids, as fatty acid esters of hydroxyl fatty acids (FAHFA) and AcylCoAs. The second category, Glycerolipids, was mainly comprised of diacylglycerols (DG) and triacylglycerols (TG). The third category, Glycerophospholipids, was comprised of lysophosphatidylcholines (LPC), phosphatidylcholines (PC), lysophosphatidylethanolamines (LPE), phosphatidylethanolamines (PE), phosphatidylglycerols (PG), Bismonoacylglycerophosphates (BMP), phosphatidylinositols (PI), phosphatidylserines (PS) and phosphatidic acids (PA). The fourth category, Sphingolipids, was comprised of ceramides (Cer), ceramide phosphates (CerP), glycosylceramides (HexCer), sulfatides and sphingomyelins (SM). The fifth category, Sterol lipids, contains only cholesteryl esters (CE).

The fatty acid profile showed marked differences between WM and GM (**Table 2**). Lower levels of 16:0 (42%), 18:0 (13%), 18:2n-6 (73%), 18:3n-3 (17%), 18:4n-3 (71%), 20:4n-6 (16%), 22:0 (70%), 22:5n-6 (49%), 22:5n-3 (68%), 22:6n-3 (75%) and 24:5n-3 (55%) whereas higher levels of 16:1n-7 (87%), 18:1n-9 (63%), 20:1n-9 (200%), 20:2n-6 (115%), 22:1n-9 (73%), 22:4n-6 (49%), 24:0 (152%), 24:1n-9 (319%), and 24:6n-3 (257%) were detected in the WM compared with the GM. Differences in FA composition results in lower levels of SFA (26%) and PUFA (31%), including PUFAn-3 (55%) and PUFAn-6 (13%), and higher levels of MUFA (67%), and ACL (1%) in WM, leading to lower values of DBI (6%) and PI (36%).

**Table 2:** Fatty acid composition, general indexes, and estimated enzyme activities in the cerebral cortex area 8 (grey matter: GM) and white matter of the *centrum semi-ovale* of the frontal lobe (WM) in middle-aged individuals without NFTs and SPs in any brain region.

|  | Grey matter<br>(GM) | White matter<br>(WM) | p-value |
| --- | --- | --- | --- |
| **14:0** | 2.51 [2.27;2.82] | 2.26 [2.06;2.40] | 0.394 |
| **16:0** | 26.0 [25.9;26.8] | 14.9 [14.8;15.3] | **0.011** |
| **16:1n-7** | 1.54 [1.46;1.64] | 2.88 [2.75;3.02] | **0.011** |
| **18:0** | 25.0 [24.6;25.2] | 21.6 [21.3;22.2] | **0.011** |
| **18:1n-9** | 20.0 [18.5;20.8] | 32.6 [31.8;33.2] | **0.011** |
| **18:1n-7** | 4.92 [4.62;5.19] | 5.31 [4.78;5.57] | 0.286 |
| **18:2n-6** | 0.96 [0.82;1.16] | 0.26 [0.25;0.33] | **0.011** |
| **18:3n-3** | 0.12 [0.11;0.13] | 0.10 [0.08;0.11] | **0.019** |
| **18:4n-3** | 1.48 [1.37;1.61] | 0.43 [0.37;0.50] | **0.011** |
| **20:0** | 0.29 [0.28;0.31] | 0.28 [0.27;0.29] | 0.522 |
| **20:1n-9** | 1.22 [1.12;1.27] | 3.76 [3.28;3.83] | **0.011** |
| **20:2n-6** | 0.48 [0.46;0.51] | 1.03 [0.86;1.11] | **0.019** |
| **20:3n-3** | 0.59 [0.56;0.62] | 0.68 [0.62;0.80] | 0.136 |
| **20:4n-6** | 4.54 [4.23;4.70] | 2.90 [2.81;2.95] | **0.011** |
| **20:3n-6** | 0.36 [0.30;0.40] | 0.25 [0.21;0.28] | 0.088 |
| **22:0** | 0.23 [0.20;0.27] | 0.07 [0.07;0.08] | **0.011** |
| **20:5n-3** | 0.87 [0.76;1.21] | 0.48 [0.46;0.52] | 0.055 |
| **22:1n-9** | 0.11 [0.10;0.12] | 0.19 [0.18;0.22] | **0.011** |
| **22:4n-6** | 2.20 [1.95;2.40] | 3.29 [3.02;3.53] | **0.011** |
| **22:5n-6** | 0.55 [0.40;0.73] | 0.28 [0.26;0.33] | **0.011** |
| **22:5n-3** | 0.28 [0.25;0.30] | 0.09 [0.07;0.09] | **0.011** |
| **24:0** | 0.40 [0.38;0.45] | 1.01 [0.95;1.04] | **0.011** |
| **22:6n-3** | 2.90 [2.79;3.18] | 0.71 [0.69;0.73] | **0.011** |
| **24:1n-9** | 0.75 [0.68;0.81] | 3.08 [2.95;3.26] | **0.011** |
| **24:5n-3** | 1.51 [1.22;1.75] | 0.68 [0.47;0.87] | **0.033** |
| **24:6n-3** | 0.14 [0.13;0.16] | 0.50 [0.43;0.54] | **0.011** |
| **SFA** | 54.7 [54.1;55.7] | 40.4 [40.0;41.5] | **0.011** |
| **UFA** | 45.3 [44.3;45.9] | 59.6 [58.5;60.0] | **0.011** |
| **PUFA** | 17.1 [16.8;17.5] | 11.6 [11.5;12.1] | **0.011** |
| **MUFA** | 28.6 [27.3;29.0] | 47.8 [46.8;48.6] | **0.011** |
| **PUFAn-3** | 8.33 [8.23;8.35] | 3.77 [3.66;4.21] | **0.011** |
| **PUFAn-6** | 9.01 [8.48;9.48] | 7.85 [7.76;8.23] | 0.136 |
| **ACL** | 17.8 [17.7;17.9] | 18.1 [18.1;18.1] | **0.011** |
| **DBI** | 101 [99.0;104] | 95.1 [94.7;95.7] | **0.011** |
| **PI (a)** | 80.9 [80.2;83.2] | 50.5 [48.6;52.6] | **0.011** |
| **PI (b)** | 56.9 [56.2;58.5] | 36.5 [35.3;37.7] | **0.011** |
| **Δ9(n-7)** | 0.06 [0.06;0.06] | 0.19 [0.18;0.20] | **0.011** |
| **Δ9(n-9)** | 0.80 [0.74;0.83] | 1.49 [1.43;1.56] | **0.011** |
| **Δ5(n-6)** | 12.6 [11.7;14.2] | 10.3 [10.2;13.1] | 0.286 |
| **Δ6(n-3)** | 12.0 [11.5;12.2] | 5.13 [3.86;5.31] | **0.011** |
| **Δ6(n-3)** | 0.09 [0.09;0.10] | 0.69 [0.62;1.03] | **0.011** |
| **Elovl3(n-9)** | 0.06 [0.06;0.06] | 0.12 [0.10;0.12] | **0.011** |
| **Elovl6** | 0.93 [0.90;0.96] | 1.45 [1.43;1.48] | **0.011** |
| **Elovl1-3-7a** | 0.01 [0.01;0.01] | 0.01 [0.01;0.01] | 0.394 |
| **Elovl1-3-7b** | 0.82 [0.74;0.89] | 0.27 [0.26;0.27] | **0.011** |
| **Elovl1-3-7c** | 1.88 [1.39;2.37] | 14.2 [11.8;16.1] | **0.011** |
| **Elovl5(n-6)** | 0.48 [0.42;0.57] | 3.72 [2.65;4.35] | **0.011** |
| **Elovl2-5 (n-6)** | 0.51 [0.48;0.52] | 1.12 [1.01;1.25] | **0.011** |
| **Elovl 2-5(n-3)** | 0.36 [0.30;0.38] | 0.17 [0.12;0.20] | 0.136 |
| **Elovl 2(n-3)** | 5.43 [4.63;6.12] | 8.95 [8.34;9.39] | **0.033** |
| **Perox β-ox** | 1.48 [1.33;1.60] | 0.22 [0.20;0.23] | **0.011** |

Values are median [Q1;Q3] from 4-6 different individuals and are expressed as mole percent. ACL: average chain length; SFA: saturated fatty acids; UFA: unsaturated fatty acids; MUFA: monounsaturated fatty acids; PUFA: polyunsaturated fatty acids; PUFAn-6: PUFA n-6 series; PUFAn-3: PUFA n-3 series; DBI: double-bond index; PI: peroxidizability index. Number of samples: GM = 4; WM = 6; p-values <0.05 highlighted in bold.

Modifications in desaturase and elongase activities could be behind these specific lipotypes. Thus, the estimation of these activities showed that the WM was characterized by a lower desaturase delta-6(n-3), elongase 1-3-7b, and peroxisomal beta-oxidation activities, whereas desaturase delta-9 and the rest of the elongases activity were significantly higher compared with the GM. 3-ketoacyl-CoA thiolase (ACAA1) immunoreactivity was found in neurons and glial cells. In neurons, moderate immunoreactivity decorated the cytoplasm; in glial cells, ACAA1 immunoreactivity formed small cytoplasmic granules. Fatty acid synthase (FAS) immunoreactivity was found in the cytoplasm of neurons and astrocytes; stearoyl-CoA desaturase (SCD) in neurons, but no immunostaining was detected in WM (**Figure 3**). Thus, enzymes linked to β-oxidation in peroxisomes (ACAA1), synthesis of palmitate from acetyl-CoA and malonyl-CoA (FAS), and integral membrane proteins implicated in fatty acid biosynthesis (SCD) are localized in neurons and glial cells.

**Figure 3:**
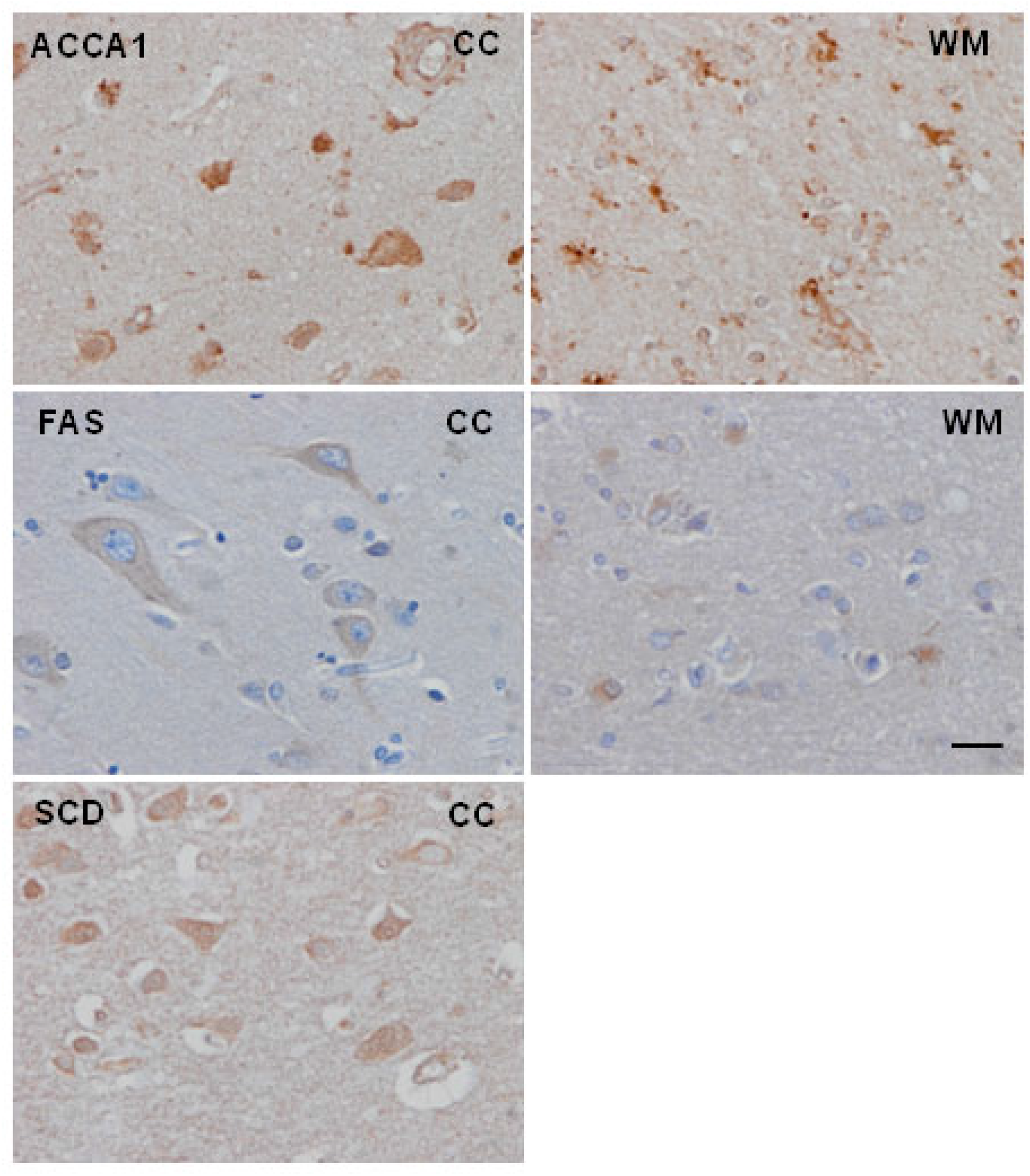
ACAA1 (3-ketoacyl-CoA thiolase) immunoreactivity was found in neurons and glial cells. In neurons, moderate immunoreactivity decorated the cytoplasm; in glial cells, ACAA1 immunoreactivity formed small cytoplasmic granules. FAS (fatty acid synthase) immunoreactivity was found in the cytoplasm of neurons and astrocytes; SCD (stearoyl-CoA desaturase) in neurons; no immunostaining was detected in the white matter. Paraffin sections slightly counterstained with haematoxylin, bar = 25μm.

### 3.2. Changes in WM and GM lipidomes associated with sAD progression

We compared the lipidome profile of GM and WM from individuals without NFT and SP pathology and AD cases grouped by their Braak stage: I-II, III-IV, and V-VI. Our results showed that diseases progression had little impact in the differences of FA profile between GM and WM. In the GM, sAD progression affected 20:4n-6 (Rho=0.439, p=0.040), 22:5n-3 (Rho= -0.693, p=0.0003), 22:6n-3 (Rho= 0.450, p= 0.035) and 24:1n-9 (Rho=-0.481, p=0.023) (**Table 3**); docosapentaenoic acid, 22:5n-3 (p=0.017), was the unique fatty acid negatively correlated with AD progression. Interestingly, AD progression was also associated with an increase in GM’s DBI (Rho=0.425, p=0.048), and PI (Rho=0.419, p=0.052), together with changes in delta-6 desaturase (Rho=0.456, p=0.032), elovl6 (Rho= - 0.587, p=0.004), elovl2-5(n-6 and n-3) (Rho= -0.426, p=0.047 and Rho= -0.519, p=0.013), respectively), elovl2 (Rho=0.537, p=0.009) activities, and peroxisomal β-oxidation (Rho=0.495, p=0.019) (**Table 3**). In contrast, reinforcing lipid resilience of WM, AD progression was accompanied by differences only in 16:1n-7 (Rho=0.405, p=0.044), 18:1n-7 (Rho=0.430, p=0.031), 22:0 (Rho=0.591, p=0.001), 22:4n-6 (Rho= -0.431, p=0.031); and DBI (Rho= -0.369, p=0.068), and PI (Rho= -0.358, p=0.078) (**Table 4**). Behenic acid, 22:0, was the only molecule that positively correlated with AD progression (Rho and p corrected value <0.05). AD progression was also associated with changes in enzymatic activities involved in fatty acid biosynthesis specifically compromising elovl6 (Rho= -0.363, p=0.073), elovl1-3-7b (Rho=0.590, p=0.001), elovl1-3-7c (Rho= -0.584, p=0.002), elovl2 (Rho= -0.387, p=0.055), and peroxisomal β-oxidation (Rho=0.402, p=0.045) (**Table 4**). Once adjusted, only elovl1-3-7b and elovl1-3-7c maintained a significant correlation with AD progression. Reinforcing the importance of anatomic location, note that the DBI and PI correlations with AD progression in GM are inverse whereas in WM are direct.

**Table 3:**
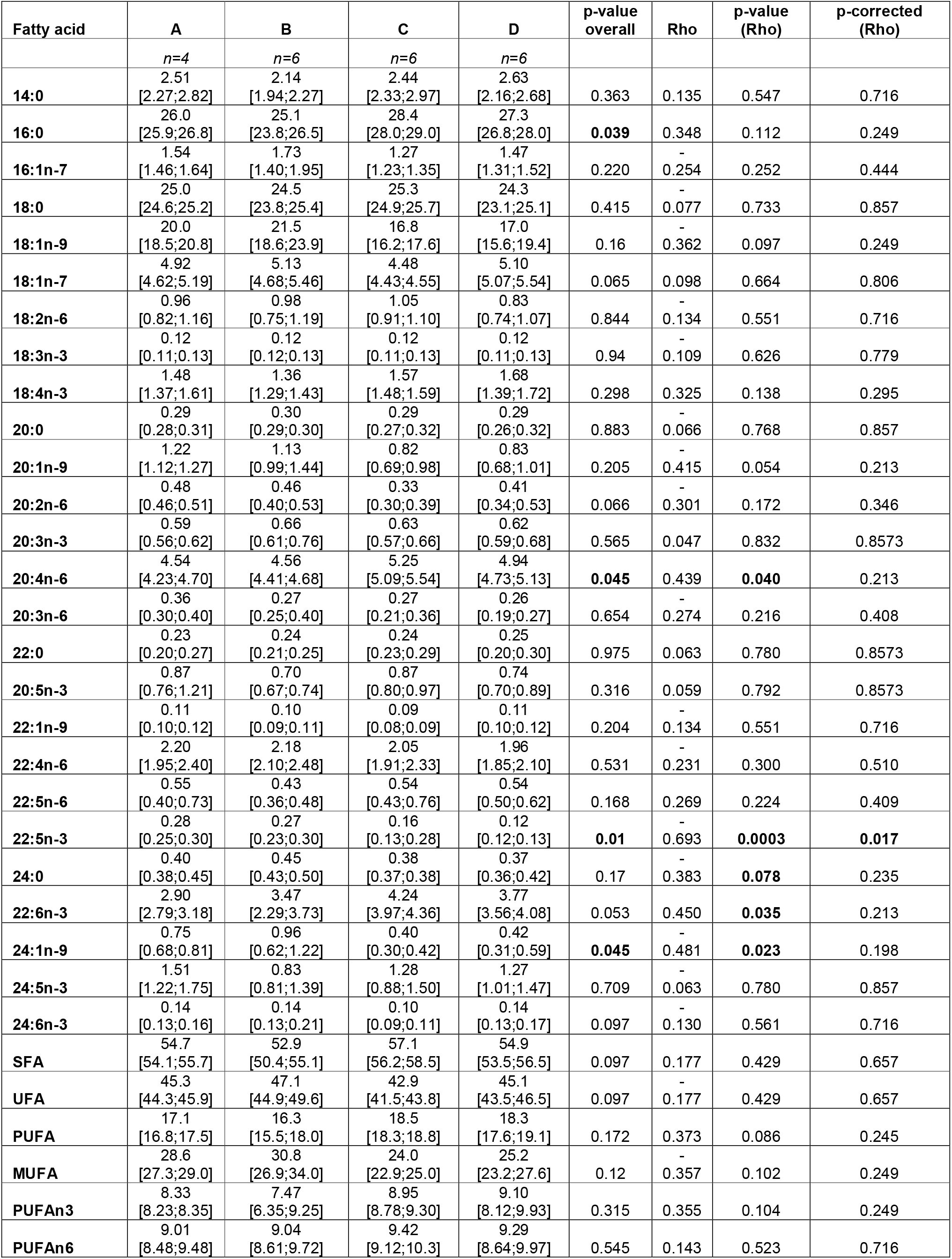
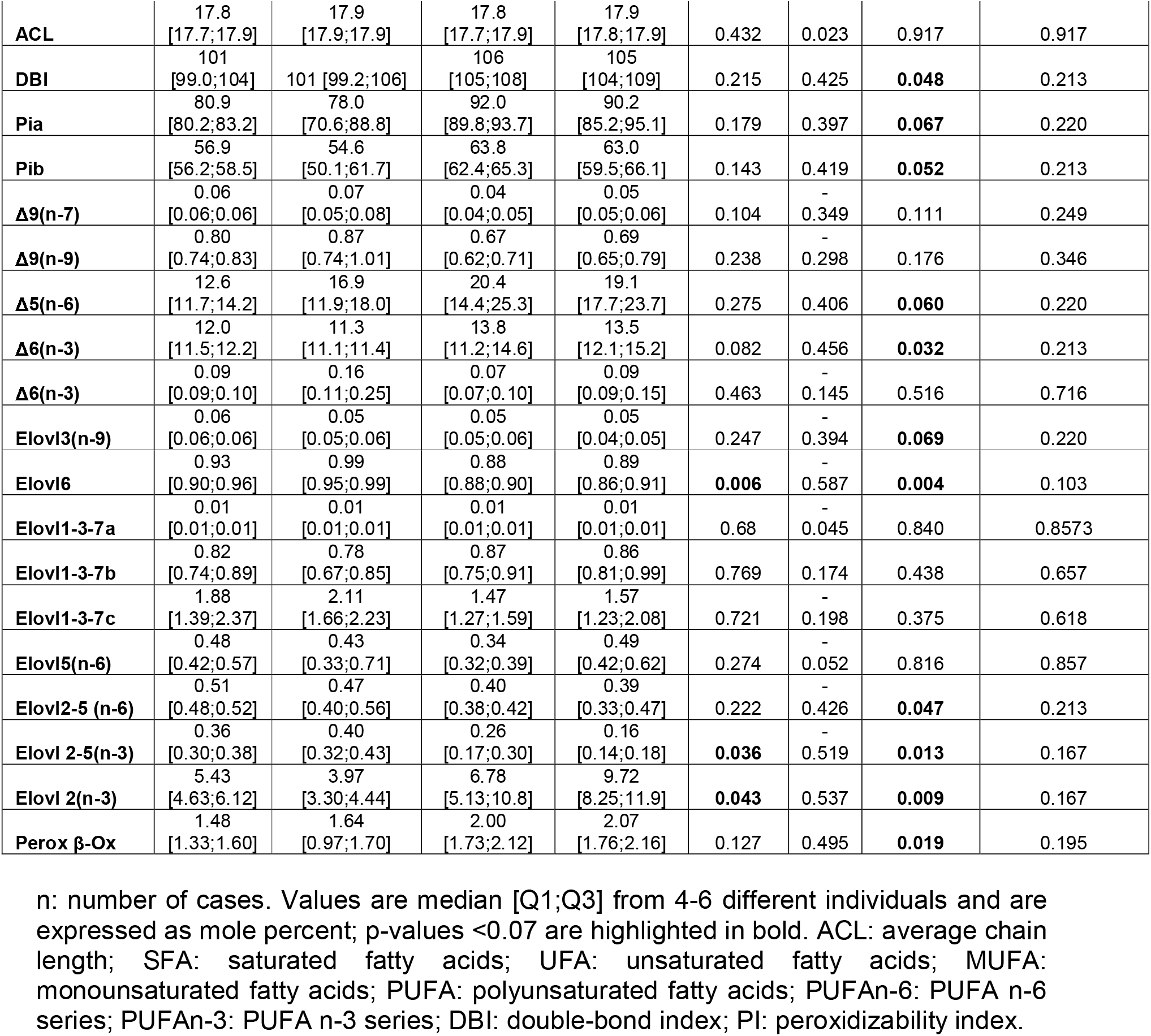
Fatty acid composition, general indexes, and estimated enzyme activities in the frontal cortex (GM) in middle-aged individuals without NFTs and SPs (A) and in cases at AD stages I-II/0-A (B), III-IV/0-B (C), and V-VI/B-C (D).

**Table 4:** Fatty acid composition, general indexes, and estimated enzyme activities in the **WM** in middle-aged individuals without NFTs and SPs (A) and in cases at AD stages I-II/0-A (B), III-IV/0-B (C), and V-VI/B-C (D).

| Fatty acid | A | B | C | D | p-value overall | Rho | p-value (Rho) | p-corrected (Rho) |
| --- | --- | --- | --- | --- | --- | --- | --- | --- |
|  | N=6 | N=7 | N=6 | N=6 |  |  |  |  |
| 14:0 | 2.26<br>[2.06;2.40] | 2.16<br>[1.93;2.36] | 1.89<br>[1.46;2.08] | 2.21<br>[2.06;2.33] | 0.175 | -0.144 | 0.491 | 0.783 |
| 16:0 | 14.9<br>[14.8;15.3] | 15.4<br>[15.2;15.6] | 16.4<br>[16.2;16.5] | 15.6<br>[15.0;16.3] | 0.194 | 0.293 | 0.153 | 0.482 |
| 16:1n-7 | 2.88<br>[2.75;3.02] | 3.04<br>[2.84;3.13] | 3.02<br>[2.96;3.07] | 3.16<br>[3.11;3.31] | 0.159 | 0.405 | **0.044** | 0.331 |
| 18:0 | 21.6<br>[21.3;22.2] | 21.3<br>[21.2;21.4] | 21.8<br>[21.2;22.0] | 21.5<br>[21.2;21.7] | 0.594 | -0.110 | 0.598 | 0.811 |
| 18:1n-9 | 32.6<br>[31.8;33.2] | 32.8<br>[32.4;33.6] | 32.9<br>[32.5;33.0] | 32.2<br>[31.9;32.4] | 0.253 | -0.159 | 0.447 | 0.735 |
| 18:1n-7 | 5.31<br>[4.78;5.57] | 5.36<br>[4.96;5.64] | 5.50<br>[5.43;5.77] | 5.86<br>[5.63;6.37] | 0.163 | 0.430 | **0.031** | 0.321 |
| 18:2n-6 | 0.26<br>[0.25;0.33] | 0.28<br>[0.24;0.42] | 0.38<br>[0.35;0.43] | 0.25<br>[0.24;0.30] | 0.236 | 0.048 | 0.819 | 0.889 |
| 18:3n-3 | 0.10<br>[0.08;0.11] | 0.12<br>[0.11;0.14] | 0.13<br>[0.11;0.14] | 0.11<br>[0.11;0.12] | 0.076 | 0.283 | 0.168 | 0.482 |
| 18:4n-3 | 0.43<br>[0.37;0.50] | 0.44<br>[0.43;0.49] | 0.47<br>[0.41;0.59] | 0.51<br>[0.49;0.54] | 0.328 | 0.366 | **0.071** | 0.331 |
| 20:0 | 0.28<br>[0.27;0.29] | 0.28<br>[0.26;0.29] | 0.26<br>[0.24;0.27] | 0.29<br>[0.28;0.31] | 0.173 | 0.087 | 0.677 | 0.825 |
| 20:1n-9 | 3.76<br>[3.28;3.83] | 3.79<br>[2.83;4.15] | 3.46<br>[3.15;3.50] | 3.36<br>[2.94;3.61] | 0.644 | -0.262 | 0.205 | 0.524 |
| 20:2n-6 | 1.03<br>[0.86;1.11] | 0.92<br>[0.88;0.94] | 0.89<br>[0.89;0.96] | 1.07<br>[0.97;1.21] | 0.129 | 0.188 | 0.367 | 0.646 |
| 20:3n-3 | 0.68<br>[0.62;0.80] | 0.63<br>[0.61;0.68] | 0.55<br>[0.54;0.68] | 0.61<br>[0.55;0.70] | 0.473 | -0.250 | 0.226 | 0.541 |
| 20:4n-6 | 2.90<br>[2.81;2.95] | 2.76<br>[2.65;2.90] | 2.95<br>[2.87;3.05] | 2.81<br>[2.51;2.89] | 0.334 | -0.050 | 0.809 | 0.889 |
| 20:3n-6 | 0.25<br>[0.21;0.28] | 0.20<br>[0.20;0.34] | 0.22<br>[0.19;0.26] | 0.21<br>[0.19;0.22] | 0.602 | -0.277 | 0.179 | 0.482 |
| 22:0 | 0.07<br>[0.07;0.08] | 0.08<br>[0.07;0.09] | 0.11<br>[0.08;0.11] | 0.11<br>[0.10;0.11] | **0.037** | 0.591 | **0.001** | **0.036** |
| 20:5n-3 | 0.48<br>[0.46;0.52] | 0.41<br>[0.40;0.44] | 0.45<br>[0.42;0.49] | 0.51<br>[0.45;0.58] | 0.367 | 0.060 | 0.774 | 0.877 |
| 22:1n-9 | 0.19<br>[0.18;0.22] | 0.18<br>[0.17;0.19] | 0.17<br>[0.16;0.18] | 0.21<br>[0.18;0.23] | 0.113 | 0.040 | 0.846 | 0.898 |
| 22:4n-6 | 3.29<br>[3.02;3.53] | 2.97<br>[2.64;3.02] | 2.88<br>[2.72;3.00] | 2.76<br>[2.59;3.04] | 0.108 | -0.431 | **0.031** | 0.321 |
| 22:5n-6 | 0.28<br>[0.26;0.33] | 0.19<br>[0.17;0.24] | 0.23<br>[0.17;0.25] | 0.22<br>[0.21;0.24] | 0.089 | -0.308 | 0.133 | 0.455 |
| 22:5n-3 | 0.09<br>[0.07;0.09] | 0.10<br>[0.07;0.11] | 0.11<br>[0.09;0.12] | 0.09<br>[0.08;0.11] | 0.434 | 0.236 | 0.254 | 0.541 |
| 24:0 | 1.01<br>[0.95;1.04] | 0.85<br>[0.81;0.97] | 0.76<br>[0.72;0.80] | 0.90<br>[0.85;0.96] | **0.041** | -0.217 | 0.296 | 0.584 |
| 22:6n-3 | 0.71<br>[0.69;0.73] | 0.77<br>[0.66;0.79] | 0.89<br>[0.86;0.95] | 0.77<br>[0.65;0.82] | **0.015** | 0.192 | 0.357 | 0.646 |
| 24:1n-9 | 3.08<br>[2.95;3.26] | 2.76<br>[2.57;3.01] | 2.75<br>[2.46;3.09] | 2.88<br>[2.75;3.73] | 0.503 | -0.035 | 0.865 | 0.900 |
| 24:5n-3 | 0.68<br>[0.47;0.87] | 0.56<br>[0.52;1.32] | 0.64<br>[0.51;1.01] | 0.54<br>[0.51;0.58] | 0.819 | -0.159 | 0.447 | 0.735 |
| 24:6n-3 | 0.50<br>[0.43;0.54] | 0.51<br>[0.39;0.54] | 0.46<br>[0.40;0.54] | 0.45<br>[0.44;0.57] | 0.951 | 0.073 | 0.725 | 0.841 |
| SFA | 40.4<br>[40.0;41.5] | 40.0<br>[39.9;40.8] | 40.8<br>[40.5;41.0] | 40.3<br>[39.8;41.6] | 0.921 | 0.082 | 0.696 | 0.825 |
| UFA | 59.6<br>[58.5;60.0] | 60.0<br>[59.2;60.1] | 59.2<br>[59.0;59.5] | 59.7<br>[58.4;60.2] | 0.921 | -0.082 | 0.696 | 0.825 |
| PUFA | 11.6<br>[11.5;12.1] | 11.2<br>[11.0;12.2] | 11.7<br>[11.4;11.9] | 11.0<br>[10.6;11.2] | 0.138 | -0.333 | 0.103 | 0.400 |
| MUFA | 47.8<br>[46.8;48.6] | 47.2<br>[46.8;48.9] | 47.6<br>[47.3;48.1] | 48.7<br>[47.3;49.5] | 0.724 | 0.138 | 0.510 | 0.788 |
| PUFAn-3 | 3.77<br>[3.66;4.21] | 3.77<br>[3.38;4.34] | 3.99<br>[3.76;4.18] | 3.60<br>[3.58;4.02] | 0.626 | -0.106 | 0.611 | 0.811 |
| PUFAn-6 | 7.85<br>[7.76;8.23] | 7.29<br>[7.02;7.86] | 7.55<br>[7.38;7.73] | 7.37<br>[7.00;7.84] | 0.457 | -0.216 | 0.297 | 0.584 |
| ACL | 18.1<br>[18.1;18.1] | 18.1<br>[18.0;18.2] | 18.1<br>[18.0;18.2] | 18.1<br>[18.0;18.1] | 0.822 | -0.104 | 0.620 | 0.811 |
| DBI | 95.1<br>[94.7;95.7] | 93.5<br>[92.0;96.5] | 94.7<br>[93.9;95.8] | 92.3<br>[91.3;93.8] | 0.102 | -0.369 | **0.068** | 0.331 |
| Pla | 50.5<br>[48.6;52.6] | 48.5<br>[46.8;52.4] | 51.2<br>[49.4;51.6] | 47.1<br>[44.7;47.9] | 0.131 | -0.327 | 0.109 | 0.400 |
| Plb | 36.5<br>[35.3;37.7] | 34.9<br>[33.9;37.8] | 36.8<br>[35.5;37.2] | 34.0<br>[32.4;34.5] | 0.135 | -0.358 | **0.078** | 0.331 |
| $\Delta 9(n-7)$ | 0.19<br>[0.18;0.20] | 0.20<br>[0.18;0.20] | 0.18<br>[0.18;0.19] | 0.21<br>[0.19;0.21] | 0.312 | 0.104 | 0.620 | 0.811 |
| $\Delta 9(n-9)$ | 1.49<br>[1.43;1.56] | 1.54<br>[1.49;1.59] | 1.50<br>[1.48;1.55] | 1.51<br>[1.50;1.52] | 0.739 | -0.022 | 0.915 | 0.934 |
| $\Delta 5(n-6)$ | 10.3<br>[10.2;13.1] | 13.2<br>[8.39;14.4] | 13.5<br>[11.8;15.8] | 13.9<br>[12.6;14.4] | 0.614 | 0.239 | 0.248 | 0.541 |
| $\Delta 6(n-3)$ | 5.13<br>[3.86;5.31] | 3.61<br>[3.39;4.12] | 3.65<br>[3.55;4.20] | 4.49<br>[4.32;4.67] | 0.186 | 0.002 | 0.989 | 0.989 |
| $\Delta 6(n-3)$ | 0.69<br>[0.62;1.03] | 0.65<br>[0.44;1.11] | 0.68<br>[0.43;1.13] | 0.94<br>[0.83;1.10] | 0.714 | 0.189 | 0.365 | 0.646 |
| Elovl3(n-9) | 0.12<br>[0.10;0.12] | 0.12<br>[0.09;0.12] | 0.11<br>[0.10;0.11] | 0.10<br>[0.09;0.11] | 0.704 | -0.237 | 0.252 | 0.541 |
| Elovl6 | 1.45<br>[1.43;1.48] | 1.39<br>[1.35;1.42] | 1.32<br>[1.30;1.36] | 1.38<br>[1.33;1.41] | 0.146 | -0.363 | **0.073** | 0.331 |
| Elovl1-3-7a | 0.01<br>[0.01;0.01] | 0.01<br>[0.01;0.01] | 0.01<br>[0.01;0.01] | 0.01<br>[0.01;0.01] | 0.097 | 0.132 | 0.528 | 0.792 |
| Elovl1-3-7b | 0.27<br>[0.26;0.27] | 0.29<br>[0.28;0.31] | 0.39<br>[0.35;0.43] | 0.34<br>[0.31;0.38] | **0.019** | 0.590 | **0.001** | **0.036** |
| Elovl1-3-7c | 14.2<br>[11.8;16.1] | 11.4<br>[10.2;11.7] | 8.07<br>[6.10;9.05] | 8.77<br>[8.20;9.78] | **0.022** | -0.584 | **0.002** | **0.036** |
| Elovl5(n-6) | 3.72<br>[2.65;4.35] | 3.20<br>[2.07;3.95] | 2.48<br>[2.14;2.54] | 4.51<br>[3.26;4.95] | 0.206 | 0.082 | 0.694 | 0.825 |
| Elovl2-5 (n-6) | 1.12<br>[1.01;1.25] | 1.08<br>[0.92;1.08] | 1.00<br>[0.92;1.02] | 1.00<br>[0.91;1.10] | 0.279 | -0.282 | 0.170 | 0.482 |
| Elovl 2-5(n-3) | 0.17<br>[0.12;0.20] | 0.19<br>[0.18;0.22] | 0.24<br>[0.21;0.27] | 0.17<br>[0.14;0.23] | 0.305 | 0.123 | 0.557 | 0.811 |
| Elovl 2(n-3) | 8.95<br>[8.34;9.39] | 5.97<br>[5.33;20.0] | 7.06<br>[6.16;8.30] | 5.98<br>[5.06;6.91] | 0.276 | -0.387 | **0.055** | 0.331 |
| Perox β-Ox | 0.22<br>[0.20;0.23] | 0.27<br>[0.23;0.28] | 0.33<br>[0.29;0.35] | 0.25<br>[0.23;0.28] | **0.023** | 0.402 | **0.045** | 0.331 |

n: number of cases. Values are median [Q1;Q3] from 4-6 different individuals and are expressed as mole percent; p-values $\leq 0.07$ are highlighted in bold. ACL: average chain length; SFA: saturated fatty acids; UFA: unsaturated fatty acids; MUFA: monounsaturated fatty acids; PUFA: polyunsaturated fatty acids; PUFAn-6: PUFA n-6 series; PUFAn-3: PUFA n-3 series; DBI: double-bond index; PI: peroxidizability index.

Multivariate and univariate statistics were performed to analyze the 2,048 features of the whole lipidome. Multivariate statistics did not show global differences in the lipidome profile GM and WM with AD progression (**Figures 4A and 4B**). For univariate statistics, we selected lipid species with a p-value < 0.05; Rho curves were applied to identify molecules associated with disease progression (**Tables 5 and 6**, and **Figures 4C and 4D**). Results showed 51 statistically different (p<0.05) lipid species in GM; 26 of them were identified based on exact mass, retention time, and or MSMS spectrum (**Table 5**). Unknown lipid species are shown in Supplementary **Table S5**. Thus, in GM AD related lipid species are grouped into four major categories: i) Fatty acyls, two esters of fatty acids, FAHFA, and one AcylCoA, Retinoyl-CoA; ii) Glycerolipids, represented by three DG, two monoacylglycerols (MG), and five TG; iii) GP, including three PC, two PE, one PG, one PI, and three PS; and iv) Sphingolipids, represented by one sulfatide and one Cer. Only 7 specific lipid species correlated with AD progression: FAHFA(26:5) (Rho = -0.480, p=0.023), TG(48:0) (Rho= - 0.429, p=0.045), TG(52:1) (Rho= -0.460, p=0.031), PC(P-41:4) (Rho= -0.619, p=0.002), PE(34:1) (Rho=0.516, p=0.013), PS(40:4) (Rho= -0.658, p=0.0008), and Cer(48:2) (Rho= 0.411, p=0.057) (**Table 5**). However, only PC(P-41:4) (p=0.027) and PS(40:4) (p=0.022) presented a Rho adjusted p value < 0.05.

**Table 5:** Significant distinctive lipidomic features in the frontal cortex area 8 in middle-aged individuals without NFTs and SPs in any brain region (A), and in cases at Braak stages I-II/0-A (B), III-IV/0-B (C), and V-VI/B-C (D).

| Class | Compound | p value | Post-hoc | Rho | p-value (Rho) | p-corr (Rho) | m/z value | RT |
| --- | --- | --- | --- | --- | --- | --- | --- | --- |
| **Fatty Acyls** |  |  |  |  |  |  |  |  |
| Fatty esters | FAHFA(24:3) a | 0.04 | A-C A-D | 0.364 | 0.095 | 0.234 | 391.2854 | 3.66 |
|  | FAHFA(26:5) a | 0.036 | A-B A-C B-C | -0.480 | **0.023** | 0.134 | 415.3025 | 3.75 |
|  | Retinoyl CoA c | 0.017 | A-B A-C B-C | -0.185 | 0.408 | 0.505 | 1072.3095 | 7.62 |
| **Glycerolipids** |  |  |  |  |  |  |  |  |
| Diradylglycerols | DG(34:3) c | 0.015 | A-B A-D B-C C-D | -0.237 | 0.288 | 0.394 | 591.4963 | 8.1 |
|  | DG(38:5) c | 0.029 | A-B A-D B-D | -0.362 | 0.097 | 0.234 | 625.5211 | 6.85 |
|  | DG(44:4) a | 0.034 | A-B A-D B-D | 0.106 | 0.637 | 0.729 | 746.7144 | 7.16 |
| Monoradylglycerols | MG(16:1) a | 0.028 | B-D C-D | 0.267 | 0.228 | 0.376 | 346.332 | 2.6 |
|  | MG(18:1) a | 0.045 | C-D | 0.271 | 0.221 | 0.376 | 374.3633 | 3.55 |
| Triradylglycerols | TG(44:3) c | 0.045 | A-C A-D C-D | 0.086 | 0.701 | 0.729 | 743.5977 | 7.92 |
|  | TG(48:0) a | 0.035 | A-B A-C B-C | -0.429 | **0.045** | 0.170 | 824.7713 | 10.14 |
|  | TG(50:0) a | 0.014 | A-B A-C B-C | -0.324 | 0.141 | 0.305 | 852.8003 | 10.31 |
|  | TG(52:1) a | 0.025 | A-B A-C B-C | -0.460 | **0.031** | 0.134 | 883.7737 | 10.33 |
|  | TG(54:4) a | 0.035 | A-B B-D | 0.240 | 0.280 | 0.394 | 900.8038 | 10.03 |
| **Glycerophospholipids** |  |  |  |  |  |  |  |  |
| Glycerophosphocholines | PC(P-41:4) a | 0.024 | A-C A-D B-C B-D | -0.619 | **0.002** | **0.027** | 880.6352 | 7.91 |
|  | PC(O-42:2)/PC(P-42:1) a | 0.035 | A-C A-D | -0.360 | 0.099 | 0.234 | 820.7033 | 7.3 |
|  | PC(40:5) a | 0.032 | A-B A-D B-D | 0.057 | 0.800 | 0.800 | 836.6162 | 7.5 |
| Glycerophosphoethanolamines | PE(34:1) a | 0.032 | B-C B-D C-D | 0.516 | **0.013** | 0.120 | 718.572 | 7.32 |
|  | PE-NMe(36:2) a | 0.032 | A-B A-D B-D | -0.258 | 0.245 | 0.376 | 756.5448 | 7.51 |
| Glycerophosphoglycerols | PG(37:0) c | 0.037 | A-B A-D C-D | 0.090 | 0.689 | 0.729 | 827.5506 | 7.31 |
| Glycerophosphoinositols | PI(34:1) a | 0.032 | A-B A-D B-D | -0.258 | 0.245 | 0.376 | 835.5253 | 6.11 |
| Glycerophosphoserines | PS(36:0) a | 0.011 | A-B A-D B-D | 0.088 | 0.695 | 0.729 | 792.5941 | 7.77 |
|  | PS(40:3) a | 0.047 | A-B A-D B-D | 0.190 | 0.396 | 0.505 | 864.5726 | 7.42 |
|  | PS(40:4) a | 0.017 | A-C A-D B-C B-D | -0.658 | **0.0008** | **0.022** | 840.5738 | 6.79 |
| **Sphingolipids** |  |  |  |  |  |  |  |  |
| Acidic glycosphingolipids | Sulfatide(43:0) a | 0.032 | A-B A-D B-D | -0.258 | 0.245 | 0.376 | 906.6894 | 8.6 |
| Ceramides | Cer(d48:2) a | 0.046 | A-D B-D | 0.411 | **0.057** | 0.186 | 746.7042 | 7.18 |

Lipidomic features with p-value <0.05 after a Kruskal-Wallis test identified by exact mass, retention time, isotopic distribution, and MS/MS spectrum; Dunn test was used for Post-hoc analysis. In bold, corrected Rho values <0.05; RT: retention time.

**Table 6:**
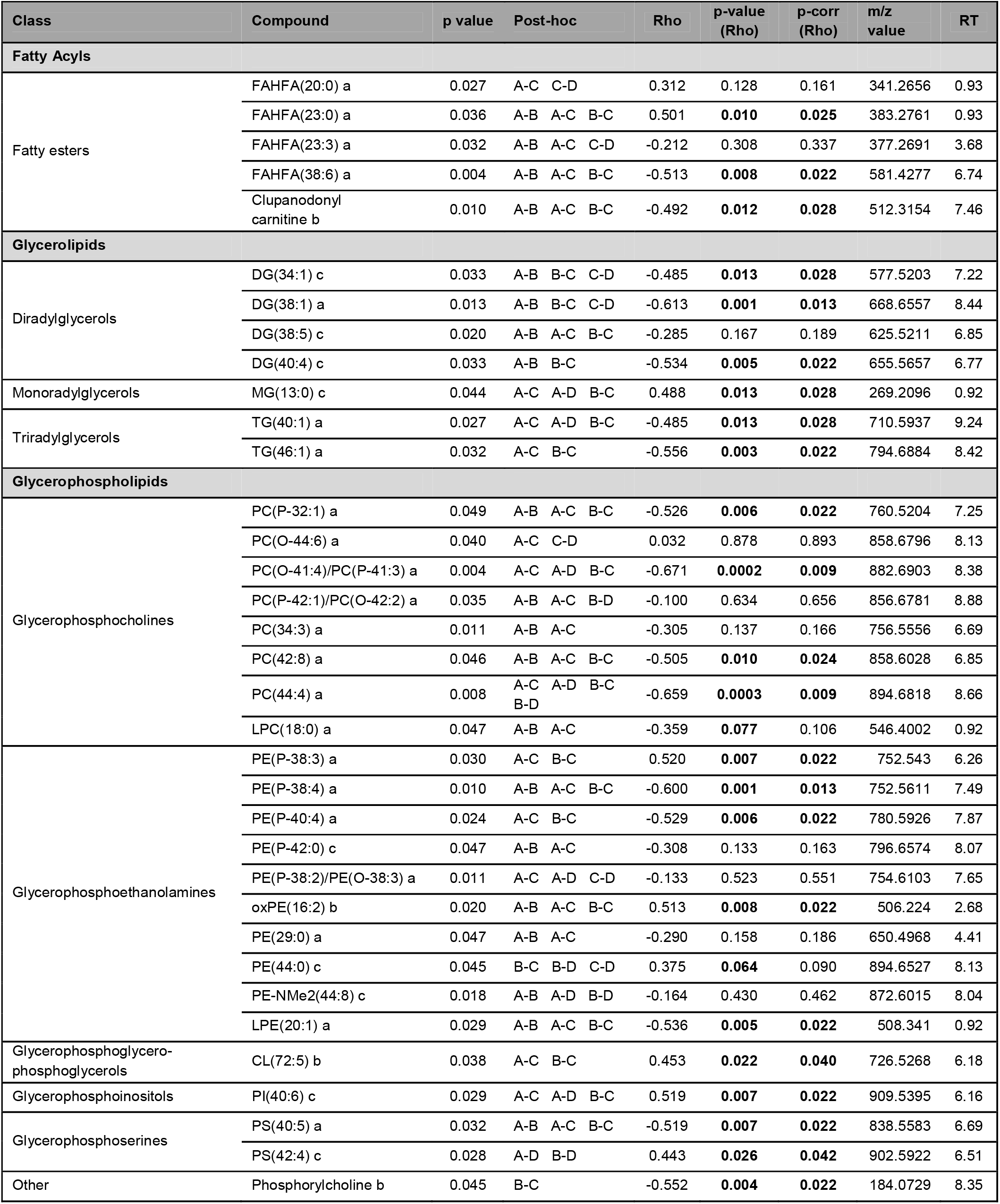
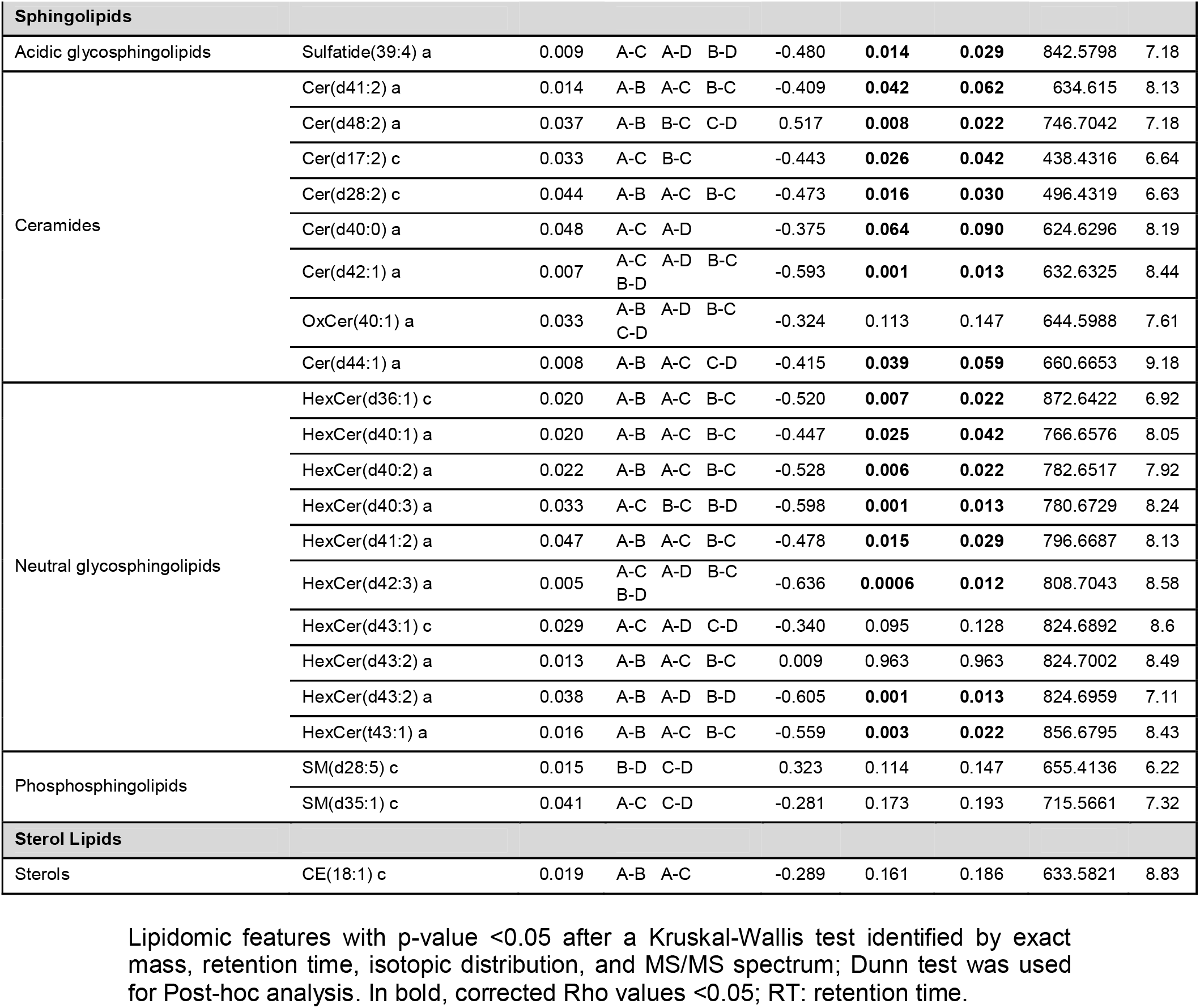
Significant distinctive lipidomic features in the white matter in middle-aged individuals without NFTs and SPs in any brain region (A), and cases at Braak stages I-II/0-A (B), III-IV/0-B (C), and V-VI/B-C (D).

**Figure 4:**
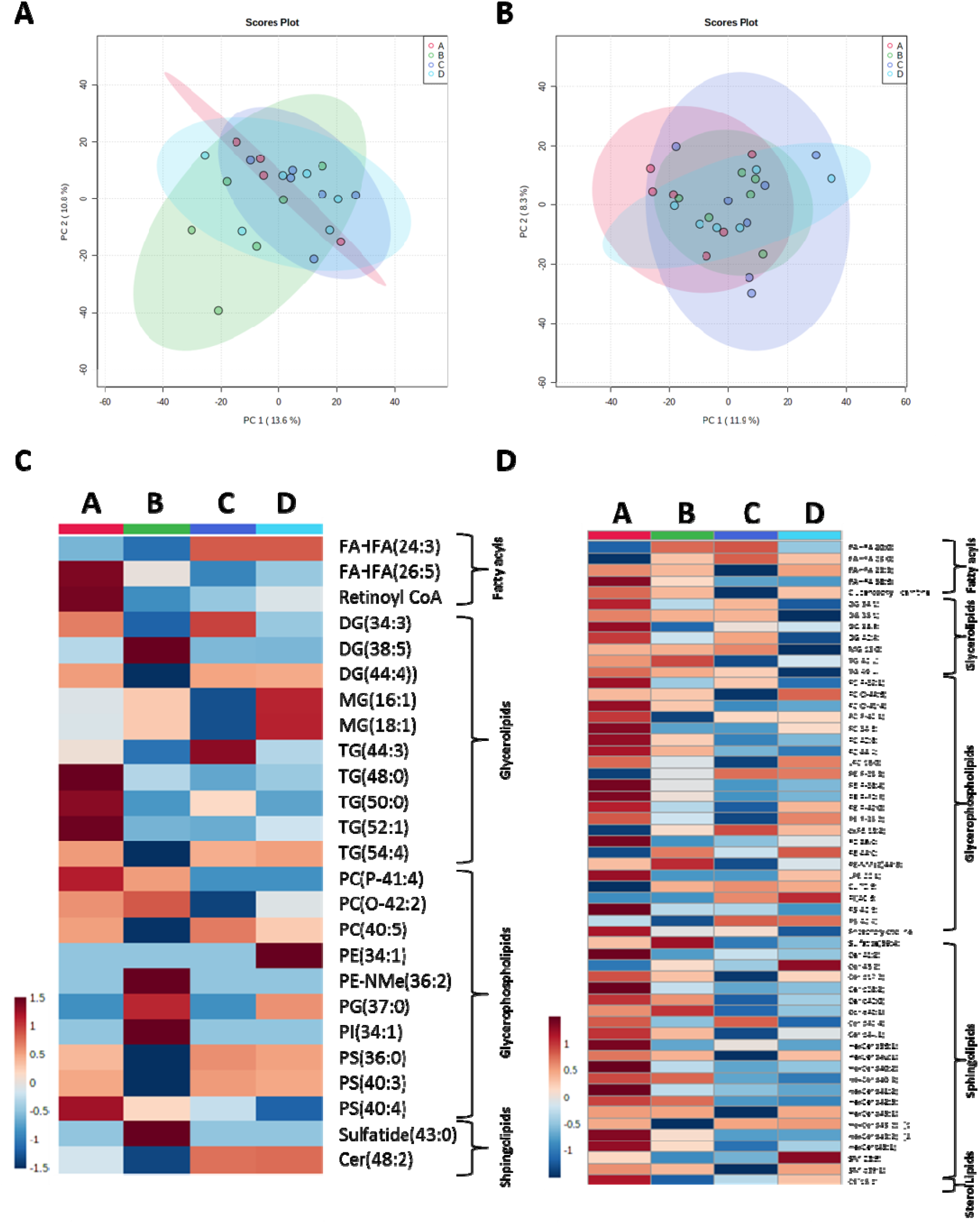
White matter is more affected by AD progression than grey matter. Principal component analyses revealed no changes associated with AD progression in grey when whole lipidome is analyzed in both grey (A) and white (B) matters. Hierarchical clustering shows the relative abundance of significantly different lipid species in grey (C) and white (D) matter. A-D in panels C and D: A: Middle-aged individuals without NFTs and SPs; B: ADI-II/0-A; C: ADIII-IV/0-B; and D: ADV-VI/B-C.

Regarding the WM, 90 lipid species were statistically different (p<0.05), and 57 of them were identified based on the criteria defined above (**Table 6** and Supplementary **Table S6** for details on unknown lipid species). Different lipid species were grouped into five major categories: i) Fatty acyls: four esters of fatty acids as FAHFAs, and one Acyl carnitine, the clupanodonyl carnitine; ii) Glycerolipids: four DG, one MG, and two TG; iii) GP: 8 PC, 10 PE, 1 CL, 1 PI, 2 PS, and the phosphorylcholine, iv) Sphingolipids, was comprised of 8 Cer, 10 HexCer, one sulfatide, and two SM; and v) Sterol lipids: one CE (**Table 6**). Changes of 41 molecules in the WM correlated with AD’s progression, 39 with a Rho adjusted p value<0.05. Among them, we found one FAHFA with a positive correlation with disease progression and another FAHFA and the clupanodonyl carnitine with a negative correlation. Three of four DG and two TG decreased, whereas MG increased with disease progression. GP identified ten molecules negatively and six positively correlated with AD progression. Remarkably, five were ether lipids, and one was an oxidized form of PE. Finally, 15 of the 16 differentially expressed sphingolipids (93.75%) -including sulfatides, ceramides, and glycosphingolipids-with Rho adjusted p-value < 0.05 correlated negatively with disease progression (**Table 6**). Only two differential lipid species were common to GM and WM, the DG(38:5) and the Cer(d48:2). These results suggest that sAD leads to more marked changes in WM than GM.

Since a relevant number of lipid species correlating with AD progression were ether lipids synthesized in peroxisomes, we analyzed the mRNA expression levels of selected genes linked to peroxisomal biogenesis, components of the peroxisomal β-oxidation, peroxisomal import of fatty acids, and fatty acyl-CoAs (see **Figure 5**). For peroxisomal biogenesis, only *PPARGC1* expression was significantly decreased in GM at advanced stages of AD (p<0.05). No differences were detected for *PPARA, PPARD*, and *PPARG*. In contrast, a gradual increase in mRNA expression of *PPARD* (p<0.05), *PPARG* (p<0.05), and *PPARGC1* (p<0.05) were identified in the WM with AD progression; *PPARA* mRNA was not altered (**Figure 5**). Regarding peroxisomal β-oxidation, only *ACAA1* mRNA expression was significantly decreased in the WM (p<0.001) but not in GM; *EHHADH* expression in GM and WM was not modified (**Figure 5**). For mRNA related to peroxisomal import of fatty acids, *ABCD2* mRNA expression was significantly decreased (p<0.05), whereas *ABCD1* and *ABCD3* mRNA were unchanged in the GM in AD. In the WM, *ABCD1* showed no changes, whereas *ABCD3* was significantly decreased (p<0.05), and ABCD2 mRNA significantly increased (p<0.05) at advanced stages of the disease (**Figure 5**).

**Figure 5:**
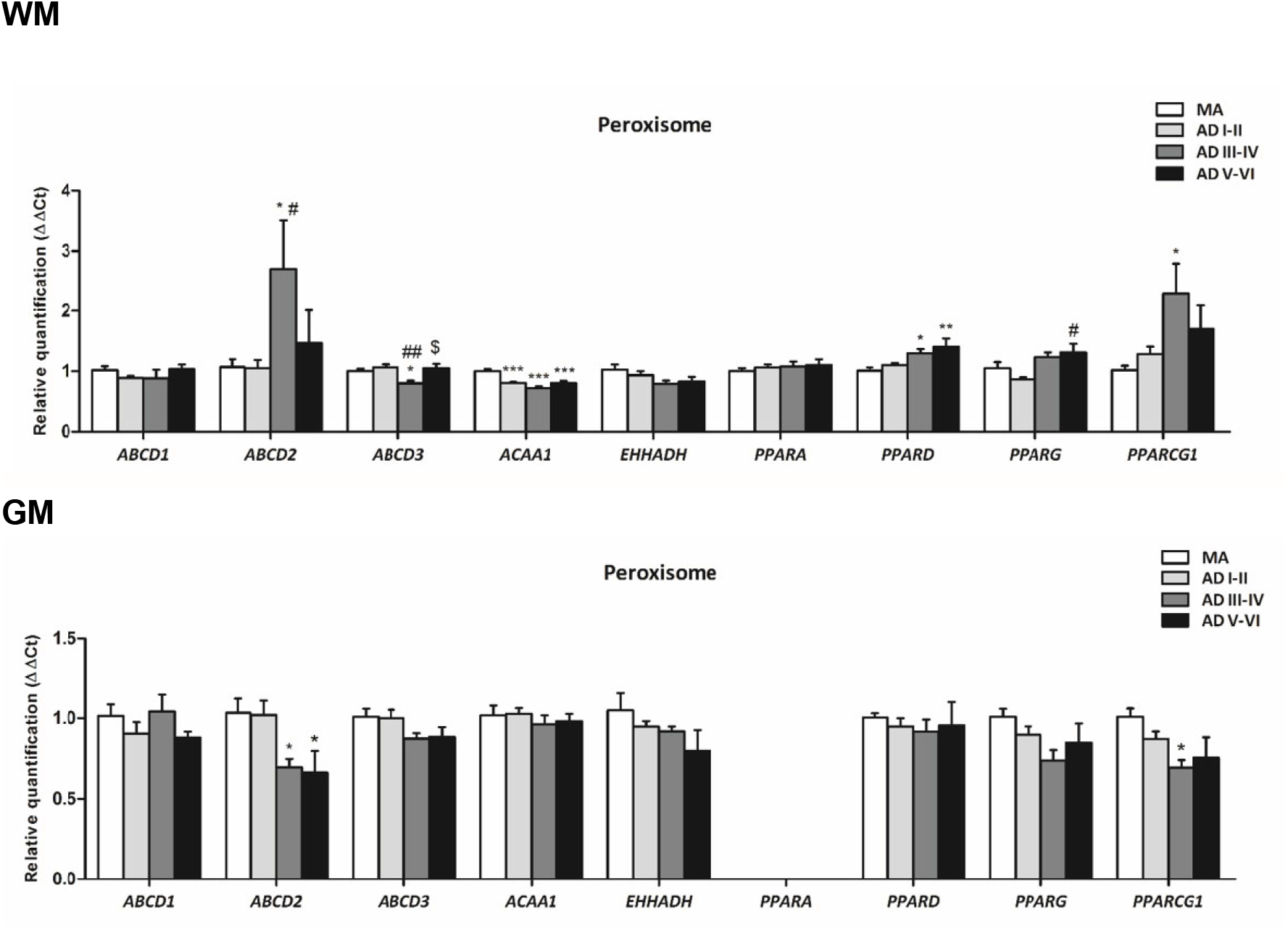
mRNA expression levels of peroxisome-related genes in the white matter of the frontal lobe (WM), and frontal cortex area 8 (GM) in middle-aged (MA) individuals without NFTs and SPs in any brain region, and at Braak stages of I-II/0-A; III-IV/0-B, and V-VI/B-C. Analyzed genes are involved in peroxisomal biogenesis: *PPARA, PPARD, PPARG*, and *PPARGC1*; peroxisomal import of fatty acids: *ABCD1, ABCD2*, and *ABCD3*; and peroxisomal β-oxidation: *ACAA1*, and *EHHADH*. T-student test or ANOVA-one way when necessary; the significance level was set at * p < 0.05, ** p < 0.01 and *** p < 0.001 vs MA; # p < 0.05 and ## p < 0.01 vs AD I-II/0-A; and $ p < 0.05 vs AD III-IV/A-B.

### 3.3. WM and GM lipidome changes linked to AD progression

Among the 25 identified lipids associated with AD progression in GM, 9 (36%) coincided with differential lipids between GM and WM. Seven of them (78%) were up-regulated in GM. Among the 57 identified lipids associated with AD progression in WM, 25 (42%) coincided with differential lipids between WM and GM. Twenty-one (83%) were up-regulated in the WM compared with GM.

## 4. Discussion

This study was designed to learn about WM and GM lipidomic modifications in middle-aged individuals without neurofibrillary tangles and senile plaques as primary markers of AD-related pathology and cases at progressive stages of sAD. Special care was taken to exclude cases with co-morbidities and associated pathologies.

In individuals with no lesions (MA), the WM is characterized by enrichment in MUFA, particularly oleic acid, 18:1n-9, and decreased content of SFA, PUFA, PUFAn-3, and PUFAn-6 and accompanied lower DBI and PI when compared with the GM. This lipid configuration is accompanied by higher delta 9-desaturase and elongase activities and decreased activity of delta-5 and delta-6 desaturases in the WM. Interestingly, both regions maintain the average chain length of 18 carbon atoms. Previous studies -covering a limited range of fatty acids-also showed similar differences in fatty acid profiles between WM and GM (O’Brien and Sampson, 1965; Sastry PS, 1985; Soderberg et al., 1991; Skinner et al., 1993). In addition, the analysis of the whole lipidome in the present study demonstrates a lower concentration of DGs, PAs, and CEs, and significant enrichment in TG, PC, PE, sulfatides, ceramides, glycosphingolipids, and sphingomyelins in the WM compared with GM. Our results are in line and go further from earlier studies showing differences in lipid composition between GM and WM (O’Brien and Sampson, 1965). Differences in the lipid composition between GM and WM have also been identified in the frontal lobe using MALDI-TOF mass spectrometry-imaging (Veloso et al., 2011) and flow infusion analysis coupled to Orbitrap™ mass spectrometry (Wood et al., 2021) in other brain regions as the temporal lobe (Man Lam et al., 2014) and the caudate nucleus (Hunter et al., 2021). Among the differential lipids in WM, we show the enrichment in lipid species belonging to the ether lipid class (alkyl and alkenyl ethers) mostly presented as TG, PC, and PE species. Considering this scenario, we demonstrate that the WM, with a higher content of UFA but with a lower degree of unsaturation, maintains the geometry and the physicochemical properties of cell membranes determining lower susceptibility to oxidative damage (lower PI), along with a higher antioxidant property linked to the high content of ether lipids. These properties constitute a more resistant condition to lipid peroxidation and, consequently, a protective environment for axonal projections.

Seminal studies reported reduced galactosylceramide (GalCer) and sulfatide and increased cholesterol and fatty acid contents in both GM and WM in AD (Yokoi et al., 1982; Soderberg et al., 1992; Svennerholm and Gottfries, 1994; Roher et al., 2002; Han et al., 2002; Hejazi et al., 2011). Levels of GalCer and sulfatide - synthesized by oligodendrocytes and major myelin components - slightly decrease in the frontal and temporal cortex and WM at stages III-IV and more markedly at stages V-VI in AD (Couttas et al., 2016). Interestingly, the activity of ceramide synthase 2, the enzyme that catalyzes the synthesis of very-long-chain ceramides, decreases in the temporal cortex at early stages and in the frontal cortex at middle-advanced stages, thus showing that alterations of ceramide synthesis occur in the early stages of AD (Couttas et al., 2016). We also observed decreased levels of specific phospholipid components of myelin in AD and reduced expression of myelin-associated proteins at advanced stages of sAD (Ferrer and Andres-Benito, 2020).

Our results show that sAD progression in the GM is associated with enrichment in PUFA, particularly 20:4n-6 and 22:6n-3, leading to a fatty acid profile with a higher peroxidizability index, consistent with previous data (Pamplona et al., 2005). In addition, the changes include a decrease in the concentration of FAHFA, TG, PC plasmalogens, and PS, together with enrichment in PE and ceramides with AD progression. Regarding the WM, increased levels of 16:1n-7, 18:1n-7, 22:0, and 22:4n-6, a fatty acid profile that determines a lower DBI and PI, and a decrease of ceramides and hexosylceramides are associated with AD progression. Moreover, the number of lipid species linked with AD progression is higher in WM than in GM. Thus, levels of 90 lipid species are statistically different comparing MA individuals and AD stages. Among 57 differential molecules with a potential identity, 41 correlated with AD progression. In contrast, in the GM, 51 lipid species showed different levels in MA and AD stages, 26 of them with a potential identity; levels of seven lipids correlate with AD progression.

To validate orthogonally these results, these AD-associated lipidome traits differences are associated with differences in the WM increased mRNA expression levels of *PPARGC1, PPARG*, and *PPARD*, linked to peroxisome biogenesis. In contrast, *PPARGC1* mRNA expression is reduced in the GM in AD. The mRNA expression of *ABCD2* and *ABCD3*, genes encoding proteins linked to the peroxisomal import of fatty acids, is also increased in the WM but not in the cerebral cortex in AD; *ABCD2* in the GM is reduced indeed. Finally, *ACAA1* mRNA expression, involved in peroxisomal β-oxidation, is reduced in the WM but preserved in the GM in sAD. These results agree with previous data suggesting the implication olf PPAR in AD pathogenesis (Wójtowicz et al., 2020).

Fatty acids are inherent components of glycerolipids, GP, and sphingolipids. The number of carbon atoms and double bonds determines the geometric traits of lipids influencing membrane organization and function (Piomelli et al., 2007). Besides, fatty acids are substrates for the generation of lipid signaling mediators, particularly relevant for PUFAn-6 and PUFAn-3 (Farooqui, 2009). An additional trait assigned to fatty acids is their chemical reactivity in oxidative conditions determining the susceptibility to oxidative damage for a given membrane (Naudi et al., 2017). Oxidant agents attack PUFA side chains much more easily than SFA and MUFA side chains (this fact is expressed by DBI and PI parameters). Minor but significant changes in the fatty acids profile mainly occur in the GM with AD progression, associated with significant vulnerability to oxidative stress conditions favored by the peroxidation-prone membrane lipid profile. By contrast, changes in fatty acid profile in the WM are associated with peroxidation-resistant membranes. Thus, we suggest that changes in GM have a clear deleterious effect, while in WM play a protective role likely as adaptative respond induced by AD pathology.

FAHFA, derived from the activity of patatin-like phospholipase domain containing 2 (PNPLA2, also known ATGL) (Patel et al., 2022) are involved in glucose homeostasis, insulin resistance and anti-inflammatory functions. FAHFA are also linked to the nuclear factor erythroid 2-related factor 2 (Nrf2), which participates in antioxidant cell defenses (Kuda et al., 2018; Wood, 2020). Of note, no previous data were available on the implication of these mediators in AD, but due to the implication of insulin resistance as a risk factor for AD (Kellar and Craft, 2020) it might be hypothesized that local FAHFA metabolism could play a pathophysiological role in AD.

DG and TG belong to the glycerolipid category. DG are cell membrane components and intermediates of lipid metabolism and act as second messengers modulating transduction proteins such as protein kinases (Carrasco and Merida, 2007; Almena and Merida, 2011; Sakane et al., 2020). Previous studies have shown altered DG levels (Chan et al., 2012; Wood et al., 2015) and deregulated protein phosphorylation (Ferrer et al., 2021) in AD. A global decrease of DG levels more marked in WM may contribute to altered membrane signaling and altered protein phosphorylation in sAD (Ferrer et al., 2021). Moreover, TG are bioenergetic compounds that, along with cholesteryl esters, are components of lipid droplets in neural cells (Teixeira et al., 2021). TG decrease in GM and WM suggests an adaptive response to higher neuronal bioenergetic demands with AD progression (Ferrer, 2009).

GP are integral components of cell membranes, substrates forming second messengers and lipid mediators, and targets and sources of oxidative stress. GP participates in a wide diversity of cell mechanisms involved in cell proliferation and differentiation, autophagy, and synthesis of other GP classes (Vance and Vance, 2004; Piomelli et al., 2007; Naudi et al., 2015). Reduced GP levels occur in aging and neurodegeneration (Kosicek and Hecimovic, 2013). Our study shows GP down-regulation, mainly affecting PC and PE levels, with AD progression, thus suggesting alterations in the architecture of neural cell membranes.

Ether lipids are a subclass of GP that show two chemical forms: alkyl ethers and alkenyl ethers or plasmalogens (Wallner and Schmitz, 2011; Dean and Lodhi, 2018). Ether lipids are primarily present as PC and PE species but have also been described as TG (Dean and Lodhi, 2018). Their biosynthesis begins in the peroxisome and is completed in the endoplasmic reticulum (Braverman and Moser, 2012; Dean and Lodhi, 2018). The physiological role of ether lipids is essentially associated with their function as membrane components with antioxidant properties (Goldfine, 2010). Lower ether lipid content is negatively associated with cancer, cardiovascular diseases, and AD (Huynh et al., 2017). The present study reveals the down-regulation of ether lipids in GM and, more markedly, in WM with AD progression. Consistent with these observations, there is an increase in oxidized PE in WM with AD progression, in agreement with enhanced lipoxidation reactions in AD (Pamplona et al., 2005).

Our transcriptomic data reveal altered expression of various genes encoding proteins linked to peroxisomes, mainly manifested by their up-regulation in the WM in AD. The contradiction between mRNA expression and down-regulation of ether lipids may result from a peroxisomal adaptation to an increased consumption rate or damage of ether lipids due to AD-induced oxidative stress conditions. However, mRNA expression levels do not necessarily parallel protein levels encoded by the corresponding genes.

Sphingolipids constitute a complex lipid group derived from N-acylsphingosine (ceramide), which is highly expressed in the human brain (Sastry, 1985). This chemical group includes a broad diversity of lipid species with structural and bioactive/messenger functions that play a vital role in the composition of lipid rafts (Posse de Chaves and Sipione, 2010; Hannun and Obeid, 2018). Sphingolipids regulate membrane physiology and cell biology (e.g., oxidative stress, apoptosis, and cell survival); they are involved in pathological conditions such as cardiovascular diseases and neurodegeneration (Trayssac et al., 2018; Meikle and Summers, 2017; Hannun and Obeid, 2018; Martin et al., 2010; Fabelo et al., 2011; Marin et al., 2017; Diaz et al., 2018). Our lipidomic study shows limited sphingolipid alterations in GM but a higher content of sphingolipids in WM in AD. In GM, only a ceramide (Cer(d48:2) correlates positively with AD progression. In contrast, 17 lipid species correlate with AD progression in WM. Only one lipid species (Cer(d48:2) positively correlates, while sixteen negatively correlate with AD progression. These results agree with recently described alterations of sphingolipid metabolism in AD, related to amyloid precursor protein-induced changes in the mitochondria-endoplasmic reticulum communications (Pera et al., 2017).

Regarding other lipid species, sulfatides, ceramides, and glycosphingolipids are decreased with AD progression, thus suggesting a negative impact on lipid raft structure and function. Lipid rafts are membrane microdomains that facilitate intercellular interactions through membrane ion channels and various signaling receptors. Membrane proteins and components of the cytoskeleton anchor in and bind to lipid rafts and regulate receptor activation, signaling pathways, membrane protein trafficking, neurotransmission, cytoskeleton, and cellular polarity (Head et al., 2014). Our findings point to alterations in lipid rafts composition since two major components, sphingolipids and phospholipids are affected in AD. Present findings are in line with previous observations showing altered lipid composition of lipid rafts in brain aging and AD (Diaz et al., 2018; Fabelo et al., 2014; Fabelo et al., 2012; Martin et al., 2010). Moreover, experimental pieces of evidence point to the facilitation of β-amyloid production resulting from abnormalities in the lipid raft composition in sAD (Diaz et al., 2015; Fabiani and Antonillini, 2019; Vetrivel and Thinakaran, 2010).

In summary, the present study characterizes the lipidome in the GM of the frontal cortex and WM of the frontal lobe *centrum semi-ovale* in middle-aged individuals and at progressive stages of AD. The WM is characterized by a fatty acid profile resistant to lipid peroxidation and enrichment of various lipid classes, mainly ether lipids, compared to GM. AD progression is associated with an altered lipidomic profile with a higher impact on WM than in GM. In particular, WM in AD is characterized by a progressive decline in the content of FAHFA, DG, TG, GP (especially ether lipids), and sphingolipids (especially sulfatides, ceramides, and glycosphingolipids). Differences among AD stages are not related to the age or gender of the individuals but the stage of the disease. Four functional categories are associated with the altered lipid classes in AD: structural membrane components, bioenergetics, antioxidant protection, and bioactive endogenous lipids. Therefore lipid metabolism can be considered key players in AD pathophysiology.

## Author Contributions

Conceptualization, M.J., I.F., and R.P.; methodology, E.O., M.J., I.F., and R.P.; software, E.O., J.S., M.J., and R.P.; formal analysis, E.O., J.S:, P.A-B., M.M-G., N.M-M., J.D.G-L., and G.P-R.; investigation, M.P-O., I.F., M.J., and R.P.; resources, G.P-R., M.P-O., I.F., M.J., R.P.; data curation, E.O., J.S., M.J., M.P-O., I.F., and R.P.; writing—original draft preparation, E.O., M.J., I.F., and R.P.; writing—review and editing, M.J., I.F., and R.P.; visualization, M.J., and R.P.; supervision, M.J., and R.P.; project administration, R.P.; funding acquisition, R.P. All authors have read and agreed upon the published version of the manuscript.

## Funding

This research was funded by the Spanish Ministry of Science, Innovation, and Universities (grant RTI2018-099200-B-I00), the Diputació de Lleida-IRBLleida (PP10605 - PIRS2021), and the Generalitat of Catalonia: Agency for Management of University and Research Grants (2017SGR696) to R.P., and Instituto de Salud Carlos III grant (PI20/00155) to M.P.O. This study was co-financed by FEDER funds from the European Union (“A way to build Europe”). IRBLleida is a CERCA Programme/Generalitat of Catalonia. Part of the study was also funded by “La Caixa” Foundation under the agreement LCF/PR/HR19/52160007: HR18-00452 to I.F.

### Institutional Review Board Statement

The study was conducted according to the guidelines of the Declaration of Helsinki, the guidelines of Spanish legislation (Real Decreto 1716/2011), and the approval of the local ethics committee (CEIC/1981).

### Informed Consent Statement

Not applicable.

### Data Availability Statement

The datasets generated and/or analyzed during the current study are available from the corresponding author upon reasonable request.

## Acknowledgments

M.J. is a ‘Serra-Hunter’ Fellow. We thank T. Yohannan for editorial help.

