## Supplemental Material for "Lipidomic alterations in the cerebral cortex and white matter in sporadic Alzheimer’s disease"

**Table S1**. Class representative and extraction internal standards added to the samples in untargeted lipidomics analysis.

| COMPOUND | SOURCE | IDENTIFIER |
| --- | --- | --- |
| 1,3(d5)-dihexadecanoyl-glycerol | Avanti Polar Lipids | 110537 |
| 1,3(d5)-dihexadecanoyl-2-octadecanoyl-glycerol | Avanti Polar Lipids | 110543 |
| 1-hexadecanoyl(d31)-2-(9Z-octadecenoyl)-sn-glycero-3-phosphate | Avanti Polar Lipids | 110920 |
| 1-hexadecanoyl(d31)-2-(9Z-octadecenoyl)-sn-glycero-3-phosphocholine | Avanti Polar Lipids | 110918 |
| 1-hexadecanoyl(d31)-2-(9Z-octadecenoyl)-sn-glycero-3-phosphoethanolamine | Avanti Polar Lipids | 110921 |
| 1-hexadecanoyl-2-(9Z-octadecenoyl)-sn-glycero-3-phospho-(1'-rac-glycerol-1',1',2',3',3'-d5) | Avanti Polar Lipids | 110899 |
| 1-hexadecanoyl(d31)-2-(9Z-octadecenoyl)-sn-glycero-3-phospho-myo-inositol | Avanti Polar Lipids | 110923 |
| 1-hexadecanoyl(d31)-2-(9Z-octadecenoyl)-sn-glycero-3-[phospho-L-serine] | Avanti Polar Lipids | 110922 |
| 26:0-d4 Lyso PC | Avanti Polar Lipids | 860389 |
| 18:1 Chol (D7) ester | Avanti Polar Lipids | 111015 |
| cholest-5-en-3ß-ol (d7) | Avanti Polar Lipids | LM-4100 |
| D-erythro-sphingosine-d7 | Avanti Polar Lipids | 860657 |
| D-erythro-sphingosine-d7-1-phosphate | Avanti Polar Lipids | 860659 |
| N-palmitoyl-d31-D-erythro-sphingosine | Avanti Polar Lipids | 868516 |
| N-palmitoyl-d31-D-erythro-sphingosylphosphorylcholine | Avanti Polar Lipids | 868584 |
| Octadecanoic acid-2,2-d2 | Sigma Aldrich | 19905-58-9 |

**Table S2**. Gene symbols and Taqman probes used for RT-qPCR.

| **Gene** | **Full name** | **Taqman probe** |
| --- | --- | --- |
| *ABCD1* | ATP Binding Cassette Subfamily D Member 1 | Hs00163610_m1 |
| *ABCD2* | ATP Binding Cassette Subfamily D Member 2 | Hs00193054_m1 |
| *ABCD3* | ATP Binding Cassette Subfamily D Member 3 | Hs00161065_m1 |
| *ACAA1* | Acetyl-CoA Acyltransferase 1 | Hs01576070_m1 |
| *EHHADH* | Enoyl-CoA Hydratase And 3-Hydroxyacyl CoA Dehydrogenase | Hs00157347_m1 |
| *GUS-β* | β-glucuronidase | Hs00939627_m1 |
| *PPARA* | Peroxisome Proliferator Activated Receptor Alpha | Hs00947539_m1 |
| *PPARD* | Peroxisome Proliferator Activated Receptor Delta | Hs00606407_m1 |
| *PPARG* | Peroxisome Proliferator Activated Receptor Gamma | Hs01115513_m1 |
| *PPARGC1A* | PPARG Coactivator 1 Alpha | Hs00173304_m1 |

**Table S3.** Identified **s**ignificant distinctive lipidomic features for white and grey matter in brain tissue.

| **Class** | **Compound** | **p value** | **Regulation**  **(WM vs Cx)** | **m/z value** | **Retention time** |
| --- | --- | --- | --- | --- | --- |
| **Fatty Acyls** |  |  |  |  |  |
| Fatty esters | FAHFA(34:1;O) c | 0.0095238 | down | 556.5378 | 8.1 |
|  | FAHFA(19:0) a | 0.0095238 | down | 327.2299 | 2.7 |
|  | FAHFA(43:4) a | 0.0095238 | down | 655.5557 | 8.2 |
|  | FAHFA(47:5) a | 0.0095238 | up | 709.6028 | 8.5 |
|  | FAHFA(45:3) a | 0.0095238 | up | 685.584 | 8.5 |
|  | FAHFA(48:5) a | 0.0095238 | up | 723.6176 | 8.7 |
|  | Retinoyl CoA c | 0.0095238 | down | 1072.3095 | 7.6 |
| **Glycerolipids** |  |  |  |  |  |
| Diradylglycerols | DG(36:4) c | 0.0057411 | down | 627.5349 | 7.3 |
|  | DG(40:5) c | 0.0057411 | down | 653.5509 | 7.5 |
|  | DG(38:7) c | 0.0057411 | down | 635.5006 | 8.5 |
|  | DG(38:6) c | 0.0089113 | down | 623.5045 | 6.7 |
|  | DG(36:4) c | 0.0089113 | down | 599.5071 | 8.9 |
|  | DG(36:1) c | 0.0095238 | up | 605.5511 | 6.8 |
|  | DG(36:2) c | 0.0095238 | up | 603.5352 | 7.3 |
|  | DG(40:4) c | 0.0095238 | down | 655.5665 | 7.6 |
|  | DG(32:2) b | 0.0095238 | down | 596.5299 | 8.1 |
|  | DG(34:3) c | 0.0095238 | down | 591.4963 | 8.1 |
|  | DG(34:1) c | 0.0095238 | up | 577.5195 | 8.1 |
|  | DG(38:1) a | 0.0095238 | up | 668.6557 | 8.4 |
|  | DG(32:1) a | 0.0095238 | down | 584.5668 | 8.5 |
|  | DG(36:3) c | 0.0095238 | down | 619.5278 | 8.5 |
|  | DG(40:6) c | 0.0095238 | up | 686.5863 | 8.5 |
| Triradylglycerols | TG(O-38:0) a | 0.0095238 | up | 670.6122 | 8.5 |
|  | TG(O-40:0) a | 0.0095238 | up | 698.6435 | 8.8 |
|  | TG(46:2) a | 0.0057411 | down | 797.6659 | 9.6 |
|  | TG(58:14) c | 0.0057411 | down | 899.6381 | 7.9 |
|  | TG(50:8) c | 0.0057411 | down | 817.6354 | 7.9 |
|  | TG(52:1) a | 0.0089113 | down | 883.7737 | 10.3 |
|  | TG(48:2) a | 0.0095238 | up | 820.7044 | 8.4 |
|  | TG(46:1) a | 0.0095238 | up | 794.6884 | 8.4 |
|  | TG(38:0) a | 0.0095238 | up | 684.6278 | 8.7 |
|  | TG(53:7) c | 0.0095238 | down | 863.6926 | 9.7 |
|  | TG(48:0) a | 0.0095238 | down | 824.7713 | 10.1 |
|  | TG(56:12) c | 0.0095238 | up | 875.6482 | 8.1 |
|  | TG(42:10) c | 0.0095238 | up | 953.6822 | 8.5 |
|  | TG(56:9) c | 0.0095238 | up | 899.7119 | 9.1 |
| **Glycerophospholipids** | |  |  |  |  |
| Glycerophosphates | PA(36:1) c | 0.0057411 | down | 703.5162 | 8.2 |
|  | PA(33:3) c | 0.0089113 | down | 679.4338 | 8.4 |
|  | PA(32:1) c | 0.0095238 | down | 647.4605 | 8.9 |
|  | PA(46:2) a | 0.0095238 | up | 841.7181 | 9 |
| Glycerophosphocholines | PC(P-38:7) a | 0.0095238 | up | 788.5444 | 6.4 |
|  | PC(P-36:1) a | 0.0095238 | up | 772.5865 | 7.2 |
|  | PC(P-34:2) a | 0.0095238 | up | 742.5749 | 7.8 |
|  | PC(P-32:1) a | 0.0095238 | down | 760.5204 | 7.2 |
|  | PC(P-36:5) a | 0.0095238 | up | 808.5385 | 7.6 |
|  | PC(O-40:0) a | 0.0095238 | up | 832.6654 | 7.8 |
|  | PC(O-40:0) a | 0.0095238 | up | 832.6674 | 8 |
|  | PC(P-36:2)/PC(O-36:3) a | 0.0089113 | down | 770.5699 | 7.5 |
|  | PC(O-36:5)/PC(P-36:4) a | 0.0095238 | up | 748.5717 | 6.7 |
|  | PC(O-42:2)/PC(P-42:1) a | 0.0095238 | up | 820.7033 | 7.3 |
|  | PC(O-34:5/PC/P-34:4) a | 0.0095238 | up | 738.5456 | 7.3 |
|  | PC(P-36:2)/PC(O-36:3) a | 0.0095238 | up | 770.6071 | 7.8 |
|  | PC(O-34:1)/PC(P-34:0) a | 0.0095238 | up | 746.6084 | 7.9 |
|  | PC(O-42:6)/PC(P-42:5) a | 0.0095238 | up | 848.697 | 8.6 |
|  | PC(O-34:1)/PC(P-34:0) c | 0.0095238 | up | 726.5809 | 6.7 |
|  | PC(O-40:4)/PC(P-40:3) a | 0.0095238 | down | 868.5965 | 7.5 |
|  | PC(O-38:2)/PC(P-38:1) a | 0.0095238 | up | 844.5979 | 7.5 |
|  | PC(P-40:2)/PC(O-40:3) c | 0.0095238 | up | 806.6416 | 7.7 |
|  | PC(O-38:1)/PC(P-38:0) c | 0.0095238 | up | 782.6402 | 8.1 |
|  | PC(O-42:2)/PC-P(42:1) c | 0.0095238 | up | 836.6894 | 8.4 |
|  | PC(O-40:1)/PC(P-40:0) c | 0.0095238 | up | 810.674 | 8.4 |
|  | PC(44:12) a | 0.0057411 | down | 922.5686 | 6.9 |
|  | PC(38:5) a | 0.0095238 | down | 764.5238 | 6.7 |
|  | PC(38:6) a | 0.0095238 | down | 806.5722 | 6.8 |
|  | PC(32:1) a | 0.0095238 | up | 732.5566 | 6.9 |
|  | PC(32:0) a | 0.0095238 | down | 734.5734 | 7.4 |
|  | PC(38:4) a | 0.0095238 | down | 810.6035 | 7.5 |
|  | PC(36:2) a | 0.0095238 | up | 786.604 | 7.5 |
|  | PC(31:0) a | 0.0095238 | up | 720.5907 | 7.9 |
|  | PC(34:0) a | 0.0095238 | down | 762.6029 | 8 |
|  | PC(38:2) a | 0.0095238 | up | 814.6329 | 8 |
|  | PC(36:1) a | 0.0095238 | up | 788.6202 | 8 |
|  | PC(44:4) a | 0.0095238 | up | 894.6818 | 8.7 |
|  | PC(38:3) c | 0.0095238 | up | 812.6167 | 7.7 |
|  | PC(36:0) c | 0.0095238 | up | 790.6559 | 7.7 |
|  | PC(36:6) a | 0.0095238 | down | 854.5205 | 7 |
|  | PC(40:2) a | 0.0095238 | up | 918.6242 | 7.1 |
|  | PC(40:7) a | 0.0095238 | up | 876.6138 | 7.1 |
|  | PC(36:5) a | 0.0095238 | down | 872.5331 | 7.2 |
|  | PC(40:3) a | 0.0095238 | up | 932.64 | 7.2 |
|  | PC(40:10) c | 0.0095238 | up | 824.531 | 7.3 |
|  | PC(42:0) c | 0.0095238 | up | 854.6993 | 8.7 |
| Glycerophosphoethanolamines | PE(P-40:6) a | 0.0089113 | down | 776.562 | 7.5 |
|  | PE(P-34:2) a | 0.0095238 | up | 700.5285 | 7.1 |
|  | PE(P-36:3) a | 0.0095238 | up | 726.544 | 7.2 |
|  | PE(P-38:4) a | 0.0095238 | up | 752.5611 | 7.5 |
|  | PE(P-34:1) a | 0.0095238 | up | 702.5467 | 7.5 |
|  | PE(P-36:2) a | 0.0095238 | up | 728.5626 | 7.6 |
|  | PE(P-38:5) a | 0.0095238 | down | 750.5606 | 7.8 |
|  | PE(P-36:1) a | 0.0095238 | up | 730.5772 | 8 |
|  | PE(P-38:2) a | 0.0095238 | up | 756.5921 | 8 |
|  | PE(P-38:1) a | 0.0095238 | up | 758.6074 | 8.4 |
|  | PE(P-40:2) a | 0.0095238 | up | 784.6224 | 8.4 |
|  | PE(P-38:3) a | 0.0095238 | down | 752.543 | 6.3 |
|  | PE(P-38:6) a | 0.0095238 | down | 746.5052 | 7 |
|  | PE(P-38:4) a | 0.0095238 | up | 750.5377 | 7.5 |
|  | PE(P-40:6) a | 0.0095238 | down | 774.5365 | 7.5 |
|  | PE(P-34:1) a | 0.0095238 | up | 700.5222 | 7.5 |
|  | PE(P-36:2) a | 0.0095238 | up | 726.5369 | 7.6 |
|  | PE(P-40:4) a | 0.0095238 | up | 778.5673 | 7.9 |
|  | PE(P-36:1) a | 0.0095238 | up | 728.5525 | 8 |
|  | PE(P-38:2) a | 0.0095238 | up | 754.5675 | 8 |
|  | PE(P-42:0) c | 0.0095238 | up | 796.6577 | 8.2 |
|  | PE(P-38:1) a | 0.0095238 | up | 756.5823 | 8.4 |
|  | PE(P-40:2) a | 0.0095238 | up | 782.5973 | 8.4 |
|  | PE(P-40:1) a | 0.0095238 | up | 784.6124 | 8.8 |
|  | PE(P-36:2)/PE(O-36:3) a | 0.0095238 | up | 726.5811 | 7.2 |
|  | PE(P-38:2)/PE(O-38:3) a | 0.0095238 | up | 754.6103 | 7.7 |
|  | PE(40:5) a | 0.0057411 | down | 794.5712 | 7.5 |
|  | PE(44:10) c | 0.0057411 | down | 870.5166 | 6.8 |
|  | PE-NMe(42:6)a | 0.0057411 | down | 832.5379 | 7.6 |
|  | PE(36:1) a | 0.0095238 | up | 746.571 | 7.2 |
|  | PE(40:6) a | 0.0095238 | down | 792.5569 | 7.2 |
|  | PE(36:2) a | 0.0095238 | up | 744.5562 | 7.3 |
|  | PE(38:4) a | 0.0095238 | down | 768.5567 | 7.3 |
|  | PE(34:2) a | 0.0095238 | up | 716.56 | 7.7 |
|  | PE(40:2) a | 0.0095238 | up | 800.6169 | 7.7 |
|  | PE(40:7) a | 0.0095238 | up | 788.5371 | 6.8 |
|  | PE(36:2) a | 0.0095238 | up | 742.5733 | 7 |
|  | PE(32:0) a | 0.0095238 | up | 672.4894 | 7 |
|  | PE(40:6) a | 0.0095238 | down | 790.5333 | 7.2 |
|  | PE(36:2) a | 0.0095238 | up | 742.5316 | 7.3 |
|  | PE(38:4) a | 0.0095238 | down | 766.5311 | 7.3 |
|  | PE(40:5) a | 0.0095238 | down | 792.5643 | 7.4 |
|  | PE(44:3) b | 0.0095238 | down | 900.5848 | 7.5 |
|  | PE-NMe(38:3) a | 0.0095238 | up | 782.5222 | 7.5 |
|  | PE(40:4) a | 0.0095238 | down | 794.5611 | 7.7 |
|  | PE(40:2) a | 0.0095238 | up | 798.6372 | 7.7 |
|  | PE(34:2) a | 0.0095238 | up | 714.5364 | 7.8 |
|  | PE(40:0) a | 0.0095238 | up | 802.5869 | 7.9 |
|  | PE(40:2) a | 0.0095238 | up | 798.638 | 7.9 |
|  | PE(38:1) a | 0.0095238 | up | 772.5768 | 8.1 |
|  | PE(36:2) a | 0.0095238 | up | 742.5653 | 8.2 |
|  | PE(44:0) c | 0.0095238 | up | 840.6849 | 8.5 |
|  | PE(46:0) b | 0.0095238 | up | 922.6838 | 8.5 |
|  | PE-NMe2(44:11) c | 0.0095238 | down | 864.5665 | 6.8 |
|  | LPE(18:0) a | 0.0057411 | down | 480.3047 | 0.9 |
|  | LPE(18:1) a | 0.0095238 | up | 480.311 | 0.9 |
|  | LPE(18:1) a | 0.0095238 | up | 478.2892 | 0.9 |
|  | LPE(22:4) a | 0.0095238 | up | 528.3035 | 2.7 |
| Glycerophosphoglycerols | PG(39:8) c | 0.0095238 | down | 822.4828 | 6 |
|  | PG(28:2) a | 0.0095238 | down | 663.4561 | 7.8 |
|  | PG(44:2) a | 0.0095238 | up | 885.667 | 8.2 |
|  | PG(44:0) c | 0.0095238 | up | 889.6632 | 8.3 |
|  | PG(44:2) a | 0.0095238 | up | 885.6969 | 8.8 |
|  | PG(44:1) c | 0.0095238 | up | 887.6475 | 7.9 |
|  | BMP(34:2) a | 0.0095238 | down | 764.5774 | 7.8 |
| Glycerophosphoglycero-phosphoglycerols | CL(78:11) a | 0.0057411 | down | 762.5007 | 6.7 |
|  | CL(78:9) b | 0.0095238 | up | 764.551 | 7.7 |
|  | CL(74:5) b | 0.0095238 | up | 740.5514 | 7.8 |
| Glycerophosphoserines | PS(P-42:7) a | 0.0057411 | down | 844.5004 | 6.7 |
|  | PS(P-40:1) a | 0.0095238 | up | 828.6028 | 7.9 |
|  | PS(42:9) c | 0.0057411 | down | 856.5023 | 6.4 |
|  | PS(44:5) c | 0.0057411 | down | 892.5978 | 7.4 |
|  | PS(40:6) a | 0.0095238 | down | 836.5444 | 6.4 |
|  | PS(36:1) a | 0.0095238 | up | 790.5607 | 6.8 |
|  | PS(40:6) a | 0.0095238 | down | 834.5214 | 6.4 |
|  | PS(36:2) a | 0.0095238 | up | 786.5197 | 6.4 |
|  | PS(40:4) a | 0.0095238 | up | 838.5484 | 6.8 |
|  | PS(46:6) c | 0.0095238 | up | 900.6142 | 6.8 |
|  | PS(40:3) a | 0.0095238 | down | 840.5656 | 6.9 |
|  | PS(40:9) c | 0.0095238 | down | 828.5043 | 7 |
|  | PS(44:4) c | 0.0095238 | up | 902.6303 | 7.1 |
|  | PS(46:2) c | 0.0095238 | up | 908.6685 | 8.3 |
| **Sphingolipids** |  |  |  |  |  |
| Acidic glycosphingolipids | Sulfatide(36:1) a | 0.0095238 | up | 806.5378 | 6 |
|  | Sulfatide(43:1) a | 0.0095238 | up | 904.611 | 6.8 |
|  | Sulfatide(42:2) a | 0.0095238 | up | 888.6163 | 6.9 |
|  | Sulfatide(42:0) a | 0.0095238 | up | 892.6085 | 7.1 |
|  | Sulfatide(43:0) a | 0.0095238 | up | 906.625 | 7.3 |
|  | Sulfatide(44:2) a | 0.0095238 | up | 916.6456 | 7.3 |
|  | Sulfatide(42:1) a | 0.0095238 | up | 890.63 | 7.3 |
|  | Sulfatide(40:4) a | 0.0095238 | down | 856.5331 | 7.5 |
|  | Sulfatide(43:1) a | 0.0095238 | up | 904.645 | 7.5 |
| Ceramides | CerP(d40:1) a | 0.0095238 | up | 684.5787 | 6.9 |
|  | OxCer(48:1) a | 0.0095238 | down | 748.7278 | 7.2 |
|  | Cer(d42:3) a | 0.0095238 | up | 646.6169 | 7.9 |
|  | Cer(d40:2) a | 0.0095238 | up | 620.5983 | 7.9 |
|  | OxCer(42:2) a | 0.0095238 | up | 630.6205 | 8 |
|  | Cer(d40:6) c | 0.0095238 | down | 612.5028 | 8.1 |
|  | Cer(d43:3) a | 0.0095238 | up | 660.6303 | 8.1 |
|  | Cer(d41:2) a | 0.0095238 | up | 634.615 | 8.1 |
|  | Cer(d38:1) a | 0.0095238 | down | 594.5836 | 8.1 |
|  | Cer(d44:3) a | 0.0095238 | up | 674.6448 | 8.3 |
|  | Cer(d42:2) a | 0.0095238 | up | 630.6214 | 8.5 |
|  | Cer(d42:4) a | 0.0095238 | up | 644.6348 | 8.7 |
|  | Cer(d40:3) a | 0.0095238 | up | 618.6196 | 8.7 |
|  | Cer(d43:4) c | 0.0095238 | up | 658.6501 | 8.9 |
|  | Cer(d44:4) a | 0.0095238 | up | 672.6275 | 8.9 |
|  | Cer(d42:1) a | 0.0095238 | up | 632.6347 | 8.9 |
|  | Cer(d16:0) c | 0.0095238 | down | 332.2617 | 2.7 |
|  | Cer(d48:2) a | 0.0095238 | down | 746.7042 | 7.2 |
|  | Cer(d36:1) a | 0.0095238 | up | 564.5301 | 7.7 |
|  | Cer(d42:3) c | 0.0095238 | up | 644.5902 | 8.2 |
|  | Cer(d42:2) a | 0.0095238 | up | 692.6111 | 8.5 |
|  | Cer(d42:2) a | 0.0095238 | up | 646.6062 | 8.5 |
|  | Cer(d40:1) c | 0.0095238 | up | 620.5902 | 8.5 |
|  | Cer(d43:3) a | 0.0095238 | up | 674.6375 | 8.9 |
|  | Cer(d42:1) c | 0.0095238 | up | 648.6215 | 8.9 |
|  | Cer(d43:1) c | 0.0095238 | up | 662.6357 | 9 |
| Neutral glycosphingolipids | HexCer(d36:2) a | 0.0095238 | up | 726.5889 | 7 |
|  | HexCer(d42:4) a | 0.0095238 | up | 806.609 | 7.9 |
|  | HexCer(d40:4) a | 0.0095238 | up | 778.6568 | 7.9 |
|  | HexCer(d42:3) a | 0.0095238 | up | 808.6695 | 7.9 |
|  | HexCer(d40:2) a | 0.0095238 | up | 782.6517 | 7.9 |
|  | HexCer(d42:2) b | 0.0095238 | up | 792.6754 | 8 |
|  | HexCer(d40:1) a | 0.0095238 | up | 766.6576 | 8.1 |
|  | HexCer(d43:3) a | 0.0095238 | up | 822.684 | 8.1 |
|  | HexCer(d41:2) a | 0.0095238 | up | 796.6687 | 8.1 |
|  | HexCer(d42:3) a | 0.0095238 | up | 808.6249 | 8.2 |
|  | HexCer(d40:3) a | 0.0095238 | up | 780.6729 | 8.2 |
|  | HexCer(d44:3) a | 0.0095238 | up | 836.6953 | 8.3 |
|  | HexCer(d43:2) a | 0.0095238 | up | 824.7002 | 8.5 |
|  | HexCer(d42:4) a | 0.0095238 | up | 806.69 | 8.2 |
|  | HexCer(d42:3) a | 0.0095238 | up | 808.7043 | 8.6 |
|  | HexCer(t40:2) a | 0.0095238 | down | 812.6734 | 6.2 |
|  | HexCer(d36:2) c | 0.0095238 | up | 886.6001 | 6.6 |
|  | HexCer(t33:3) a | 0.0095238 | up | 712.5196 | 7.4 |
|  | HexCer(d42:0) a | 0.0095238 | up | 812.6537 | 7.9 |
|  | HexCer(d43:1) a | 0.0095238 | up | 824.6537 | 7.9 |
|  | HexCer(d42:2) a | 0.0095238 | up | 808.6597 | 8.1 |
|  | HexCer(t42:3) a | 0.0095238 | up | 838.6688 | 8.1 |
|  | HexCer(d42:0) a | 0.0095238 | up | 812.6538 | 8.1 |
|  | HexCer(t43:3) a | 0.0095238 | up | 852.6833 | 8.3 |
|  | HexCer(d43:0) a | 0.0095238 | up | 826.6701 | 8.3 |
|  | HexCer(d43:1) c | 0.0095238 | up | 824.6892 | 8.6 |
| Phosphosphingolipids | SM(d34:1) a | 0.0095238 | up | 703.5768 | 6.7 |
|  | SM(d36:2) a | 0.0095238 | up | 729.5938 | 6.8 |
|  | SM(d35:1) a | 0.0095238 | up | 717.5907 | 7 |
|  | SM(d40:2) a | 0.0095238 | up | 785.6552 | 7.9 |
|  | SM(d38:1) a | 0.0095238 | down | 759.6399 | 7.8 |
|  | SM(d43:2) a | 0.0095238 | up | 827.7037 | 8.7 |
|  | SM(d41:1) a | 0.0095238 | up | 801.685 | 8.7 |
|  | SM(d42:1) a | 0.0095238 | up | 815.702 | 9.1 |
|  | SM(d43:2) a | 0.0095238 | up | 871.6536 | 8 |
|  | SM(d43:2) a | 0.0095238 | up | 871.6809 | 8.5 |
| **Sterol Lipids** |  |  |  |  |  |
| Sterols | CE(15:0) c | 0.0095238 | down | 645.5232 | 8.6 |
|  | OxCE(18:1) c | 0.0095238 | down | 687.5694 | 9 |

*Lipidomic features with p-value < 0.01 (FDR < 0.07) after a Wilcoxon test. We have classified the lipid species according to the confidence of the identification: a) exact mass, retention time and MS/MS spectrum (high reliability), b) exact mass, retention time and MS/MS spectrum (medium reliability), c) exact mass and retention time*

**Table S4**. Unidentified significant distinctive lipidomic features for white and grey matter in brain tissue.

| **Class** | **Compound** |  |  | **P value** | **Regulation**  **(WM vs Cx)** | **m/z value** | **Retention time** |
| --- | --- | --- | --- | --- | --- | --- | --- |
| **Unknown** |  |  |  |  |  |  |  |
| Unknown | 398.19_0.86392 |  |  | 0.0057411 | down | 399.1973 | 0.9 |
|  | 677.5246_8.0952 |  |  | 0.0057411 | down | 678.5319 | 8.1 |
|  | 668.5212_8.6017 |  |  | 0.0057411 | down | 669.5285 | 8.6 |
|  | 280.2378_3.0610 |  |  | 0.0057411 | down | 279.2305 | 3.1 |
|  | 565.6891_7.1682 |  |  | 0.0089113 | down | 566.6964 | 7.2 |
|  | 634.5336_7.4776 |  |  | 0.0089113 | down | 635.5409 | 7.5 |
|  | 238.1211_0.8675 |  |  | 0.0095238 | down | 239.1284 | 0.9 |
|  | 796.1554_5.6956 |  |  | 0.0095238 | down | 797.1627 | 5.7 |
|  | 780.1825_5.9419 |  |  | 0.0095238 | down | 781.1898 | 5.9 |
|  | 560.5174_7.5109 |  |  | 0.0095238 | up | 561.5247 | 7.5 |
|  | 514.4072_7.5952 |  |  | 0.0095238 | down | 515.4145 | 7.6 |
|  | 527.4968_7.6575 |  |  | 0.0095238 | down | 528.5041 | 7.7 |
|  | 735.5375_7.7941 |  |  | 0.0095238 | down | 736.5448 | 7.8 |
|  | 541.5115_7.8739 |  |  | 0.0095238 | down | 542.5188 | 7.9 |
|  | 317.2983_8.0923 |  |  | 0.0095238 | down | 318.3056 | 8.1 |
|  | 492.4916_8.0789 |  |  | 0.0095238 | down | 493.4989 | 8.1 |
|  | 312.2668_8.1176 |  |  | 0.0095238 | up | 313.2741 | 8.1 |
|  | 338.2824_8.1365 |  |  | 0.0095238 | up | 339.2897 | 8.1 |
|  | 1314.2874_8.382 |  |  | 0.0095238 | up | 1315.2947 | 8.4 |
|  | 1242.2812_8.378 |  |  | 0.0095238 | up | 1243.2885 | 8.4 |
|  | 340.298_8.50841 |  |  | 0.0095238 | up | 341.3053 | 8.5 |
|  | 338.2826_8.5110 |  |  | 0.0095238 | up | 339.2899 | 8.5 |
|  | 872.2598_8.8562 |  |  | 0.0095238 | down | 873.2671 | 8.9 |
|  | 242.1491_0.9088 |  |  | 0.0095238 | down | 241.1418 | 0.9 |
|  | 310.2843_4.5641 |  |  | 0.0095238 | up | 309.2770 | 4.6 |
|  | 366.3459_6.2096 |  |  | 0.0095238 | up | 365.3386 | 6.2 |
|  | 394.3772_6.9073 |  |  | 0.0095238 | up | 393.3699 | 6.9 |
|  | 1164.2555_7.911 |  |  | 0.0095238 | down | 1163.2482 | 7.9 |
|  | 1238.2736_8.184 |  |  | 0.0095238 | down | 1237.2663 | 8.2 |
|  | 823.6816_8.2384 |  |  | 0.0095238 | up | 822.6743 | 8.2 |
|  | 633.5969_8.3553 |  |  | 0.0095238 | up | 632.5896 | 8.4 |
|  | 1386.309_8.5942 |  |  | 0.0095238 | down | 1385.3017 | 8.6 |
|  | 661.6291_8.7003 |  |  | 0.0095238 | up | 660.6218 | 8.7 |

*Lipidomic features with p-value < 0.01 (FDR < 0.07) after a Wilcoxon test non identified by MS, RT and MS/MS spectra classified as Unknowns.*

**Table S5**. Unidentified significant distinctive lipidomic features in grey matter during AD progression.

| **Class** | **Compound** |  | **p value** | **Post-hoc** | **Rho** |  | **m/z value** | **Retention time** |
| --- | --- | --- | --- | --- | --- | --- | --- | --- |
| **Unknown** | |  |  |  |  |  |  |  |
| Unknown | 1077.2719_7.635 |  | 0.010448 | A-B A-C A-D B-C |  |  | 1078.2792 | 7.64 |
|  | 1080.2548_7.635 |  | 0.031718 | A-C B-C |  |  | 1081.2621 | 7.64 |
|  | 1092.2303_7.625 |  | 0.021924 | A-B A-D B-D |  |  | 1093.2376 | 7.63 |
|  | 110.0361_0.8535 |  | 0.016152 | A-C A-D C-D |  |  | 111.0434 | 0.85 |
|  | 128.046_0.79454 |  | 0.02328 | A-C B-D C-D |  |  | 127.0387 | 0.79 |
|  | 211.1594_0.9228 |  | 0.038388 | C-D |  |  | 212.1667 | 0.92 |
|  | 214.0206_0.7680 |  | 0.024433 | A-B A-D B-D |  |  | 213.0133 | 0.77 |
|  | 239.2629_10.549 |  | 0.020236 | A-B A-C B-C |  |  | 240.2702 | 10.55 |
|  | 302.1642_2.2336 |  | 0.029211 | A-B B-D |  |  | 301.1569 | 2.23 |
|  | 308.2141_4.6456 |  | 0.048831 | A-B B-C |  |  | 307.2068 | 4.65 |
|  | 329.3286_3.2113 |  | 0.009155 | A-B A-D B-C C-D |  |  | 330.3359 | 3.21 |
|  | 335.3168_0.9753 |  | 0.028794 | A-B A-C B-C |  |  | 336.3241 | 0.98 |
|  | 340.093_0.84219 |  | 0.020709 | A-B A-D B-D |  |  | 341.1003 | 0.84 |
|  | 340.1492_0.8236 |  | 0.043647 | A-B A-C B-C |  |  | 341.1565 | 0.82 |
|  | 346.2181_7.5950 |  | 0.04852 | A-D B-C B-D |  |  | 347.2254 | 7.6 |
|  | 360.1745_0.8947 |  | 0.022657 | A-C A-D C-D |  |  | 359.1672 | 0.89 |
|  | 360.3367_0.9784 |  | 0.043616 | A-D B-D |  |  | 361.344 | 0.98 |
|  | 398.19_0.86392 |  | 0.029269 | A-B A-C B-C |  |  | 399.1973 | 0.86 |
|  | 418.309_5.52832 |  | 0.017704 | B-D C-D |  |  | 419.3163 | 5.53 |
|  | 481.3119_3.4994 |  | 0.031466 | A-C A-D C-D |  |  | 480.3046 | 3.5 |
|  | 60.0204_0.86653 |  | 0.023184 | A-B A-C |  |  | 59.0131 | 0.87 |
|  | 612.1748_7.7724 |  | 0.04124 | A-B A-C B-C |  |  | 613.1821 | 7.77 |
|  | 851.9602_7.1495 |  | 0.045484 | A-D |  |  | 850.9529 | 7.15 |
|  | 965.22_9.025290 |  | 0.016541 | A-B A-D B-C |  |  | 966.2273 | 9.03 |
|  | 997.283_7.26389 |  | 0.04957 | A-B B-D |  |  | 998.2903 | 7.26 |

*Lipidomic features with p-value < 0.05 (FDR > 0.05) after a Kruskal-Wallis’s test non identified by MS, RT and MS/MS spectra classified as Unknowns.*

**Table S6**. Unidentified significant distinctive lipidomic features in white matter during AD progression.

| **Class** | **Compound** |  |  | **p value** | **Post-hoc** | **m/z value** | **Retention time** |
| --- | --- | --- | --- | --- | --- | --- | --- |
| **Unknown** |  |  |  |  |  |  |  |
| Unknown | 1053.2917_9.502 |  |  | 0.033862 | A-B A-C B-C | 1054.299 | 9.5 |
|  | 1094.247_7.9067 |  |  | 0.022435 | B-C B-D | 1095.2543 | 7.91 |
|  | 124.0514_0.8604 |  |  | 0.040452 | A-C A-D C-D | 123.0441 | 0.86 |
|  | 126.0676_0.8716 |  |  | 0.035964 | A-C A-D C-D | 127.0749 | 0.87 |
|  | 166.1354_0.9250 |  |  | 0.025681 | A-B B-D C-D | 167.1427 | 0.93 |
|  | 172.1447_0.9338 |  |  | 0.045498 | A-B A-D B-D | 171.1374 | 0.93 |
|  | 199.1934_1.8807 |  |  | 0.037332 | A-C B-C | 200.2007 | 1.88 |
|  | 208.1917_0.9568 |  |  | 0.020253 | A-B A-D B-D | 209.199 | 0.96 |
|  | 210.1597_0.9174 |  |  | 0.023644 | B-C B-D | 209.1524 | 0.92 |
|  | 238.2174_10.554 |  |  | 0.0065421 | A-B B-C C-D | 239.2247 | 10.55 |
|  | 240.1641_0.9626 |  |  | 0.0066994 | B-C B-D C-D | 239.1568 | 0.96 |
|  | 242.1155_0.8251 |  |  | 0.031692 | A-B A-D B-C | 243.1228 | 0.83 |
|  | 250.1576_0.9329 |  |  | 0.014293 | A-B A-C B-C | 251.1649 | 0.93 |
|  | 254.1886_0.9288 |  |  | 0.026092 | A-B A-C | 255.1959 | 0.93 |
|  | 286.0828_0.8305 |  |  | 0.039072 | A-C B-C | 287.0901 | 0.83 |
|  | 324.154_0.86864 |  |  | 0.038458 | A-B A-D B-C | 325.1613 | 0.87 |
|  | 360.3337_0.9602 |  |  | 0.045854 | B-C C-D | 359.3264 | 0.96 |
|  | 388.2972_8.4307 |  |  | 0.01834 | A-B A-C B-C | 389.3045 | 8.43 |
|  | 406.2708_0.9220 |  |  | 0.025195 | A-B A-C C-D | 405.2635 | 0.92 |
|  | 446.2244_6.7663 |  |  | 0.005119 | A-B A-D B-D | 445.2171 | 6.77 |
|  | 552.3831_6.3107 |  |  | 0.028744 | A-B A-C B-C | 553.3904 | 6.31 |
|  | 592.5909_8.4160 |  |  | 0.0098526 | A-B A-C B-C | 593.5982 | 8.42 |
|  | 657.7232_7.1611 |  |  | 0.047273 | A-C A-D C-D | 656.7159 | 7.16 |
|  | 667.0593_0.9134 |  |  | 0.0038145 | A-B A-C A-D B-C C-D | 666.052 | 0.91 |
|  | 685.5312_0.9721 |  |  | 0.032858 | A-B A-C B-D | 686.5385 | 0.97 |
|  | 706.164_5.34443 |  |  | 0.040145 | A-D B-C C-D | 707.1713 | 5.34 |
|  | 720.6389_7.5765 |  |  | 0.030618 | A-C B-C | 721.6462 | 7.58 |
|  | 744.2015_0.9867 |  |  | 0.041729 | B-C B-D | 743.1942 | 0.99 |
|  | 762.1933_6.9904 |  |  | 0.039731 | A-B A-C B-D C-D | 763.2006 | 6.99 |
|  | 771.2663_8.8725 |  |  | 0.033477 | A-B A-C | 772.2736 | 8.87 |
|  | 822.2932_5.4247 |  |  | 0.049824 | A-B A-C | 823.3005 | 5.42 |
|  | 845.6082_0.9154 |  |  | 0.011592 | A-C B-D C-D | 846.6155 | 0.92 |
|  | 854.2024_6.4596 |  |  | 0.035044 | A-C A-D B-C | 855.2097 | 6.46 |

*Lipidomic features with p-value < 0.05 (FDR > 0.05) after a Kruskal-Wallis’s test non identified by MS, RT and MS/MS spectra classified as Unknowns.*
